# Mobile Human Brain Imaging using Functional Ultrasound

**DOI:** 10.1101/2024.12.08.627337

**Authors:** Sadaf Soloukey, Luuk Verhoef, Frits Mastik, Michael Brown, Geert Springeling, Bastian S. Generowicz, Djaina D. Satoer, Clemens M.F. Dirven, Marion Smits, Borbála Hunyadi, Sebastiaan K.E. Koekkoek, Arnaud J.P.E. Vincent, Chris I. De Zeeuw, Pieter Kruizinga

## Abstract

Imagine being able to study the human brain in real-world scenarios while the subject displays natural behaviors such as locomotion, social interaction, or spatial navigation. The advent of ultrafast ultrasound imaging brings us closer to this goal with functional Ultrasound imaging (fUSi), a new mobile neuroimaging technique. Here, we present real-time fUSi monitoring of brain activity during walking in a subject with a clinically approved sonolucent skull implant. Our approach utilizes personalized 3D-printed fUSi-helmets for stability, optical tracking for cross-modal validation with fMRI, advanced signal processing to estimate hemodynamic responses and facial tracking of a lick licking paradigm. These combined efforts allowed us to show consistent fUSi signals over 20 months, even during high-motion activities like walking. These results demonstrate the feasibility of fUSi for monitoring brain activity in real-world contexts, marking an important milestone for fUSi-based insights in clinical and neuroscientific research.

## Introduction

Being able to study the human brain in real-world scenarios while the subject is displaying natural behaviors such as locomotion, social interaction or spatial navigation, could be immensely valuable for our clinical and neuroscientific understanding of human brain activity^1,2^. Unfortunately, functional brain imaging techniques come with technical trade-offs which - until now - have not been able to facilitate high-quality imaging of human brain activity in such ecological contexts. Non-invasive, wearable techniques such as electroencephalography (EEG)^3^ or functional near-infrared spectroscopy (fNIRS)^4,5^ do facilitate mobility, but without allowing for high-resolution imaging of deeper brain structures. Optical techniques such as diffuse optical tomography (DOT)^5,6^ allow for non-invasive imaging with a large field of view but have low resolution and limited penetrative depth. Invasive techniques such as electrocorticography (ECoG)^7^ or intracranial EEG (iEEG)^8^ allow for high-resolution mapping of the brain, but require implantation beneath the skull, with limited implant lifetime. Finally, non-invasive, whole-brain techniques such as magnetoencephalography (MEG)^9^ and especially functional Magnetic Resonance Imaging (fMRI)^10^ dominate the current human functional brain imaging landscape, but require very large and expensive machinery, while severely restricting movement of subjects during imaging. In fact, many functional paradigms such as locomotion or speech are performed in an ‘imagined’ fashion, where the subject is asked to imagine performing the functional task instead^11–13^.

With the advent of ultrafast Doppler ultrasound imaging, a new brain imaging technique called functional Ultrasound imaging (fUSi) has emerged for neuroscientific^14,15^ and clinical use^16–18^. fUSi exploits a high-frame-rate (HFR) acquisition scheme using unfocussed transmissions to boost the sensitivity of conventional Doppler ultrasound^14^. This gain in sensitivity allows for detection of subtle changes in hemodynamics in the brain’s (micro)vasculature, which through the principle of neurovascular coupling (NVC), serve as a proxy for functional brain activity^14,19,20^. What makes fUSi unique, is its ability to combine high-resolution functional brain imaging with a high level of portability and flexibility^16,18^. Simultaneously, fUSi is contrast-agent-free and comes with all the other well-known benefits of conventional ultrasound: it is real-time, non-invasive, easy to use and comparatively cost-effective. None of the currently available brain imaging techniques are able to combine all these beneficial characteristics at once^15,17^.

Given these benefits, fUSi has found its way from a pre-clinical context to in-human, neurosurgical applications in less than a decade^16,18,21^. During awake brain surgeries for tumor resections, fUSi has been applied successfully to map out hemodynamics-based functional brain activity at a mesoscopic scale while patients performed language, motor and sensory tasks^16,18,22^. So far, the neurosurgical context has not just been an interesting use-case for fUSi, it is a necessary one as well. Given the significant attenuation and aberration of ultrasound signal through human skull bone^23^, a craniotomy is a necessary acoustic window to the human brain, making fUSi is not applicable outside of the operating room as of yet. As a consequence, fUSi is confined to the very limited context of awake brain surgeries, where time is limited, and patients are physically restricted^17^.

Clinical practice and scientific literature demonstrate other approaches to still achieve acoustic access to the brain *outside* of the operating room. Clinical use of burrholes or cranioplasties also facilitate – as a byproduct – artificial acoustic windows to the human brain^24^. Cranioplasties are performed in clinical contexts for a range of etiologies where a skull bone defect (SBD) cannot be covered with the patient’s autologous skull bone flap, such as after traumatic brain injury of bone flap infection after intracranial tumor surgery^25^. The SBD can then be covered by an artificial, patient specific implant made of materials such as Titanium Mesh (TM), Polymethyl Methacrylate (PMMA) or polyetheretherketone (PEEK)^26^. The latter two types of plastic are far more homogeneous and present with much less acoustic signal attenuation as compared to human skull bone^27^, making them sonoluscent. A handful of studies have demonstrated the potential of PMMA and PEEK for conventional B-mode ultrasound-imaging, e.g. in the context of bedside monitoring of ventricular size in neurotrauma patients^28–32^.

Recently, Rabut et al.^33^ published the first demonstration of in-human fUSi through a custom, thinned-out PMMA window, implanted in a human subject with an SBD after trauma. The authors show the feasibility of fUSi for mapping and decoding of task-modulated cortical activity through PMMA during functional tasks involving gaming and guitar playing^33^.

In this paper we show the next step: successful use of fUSi in a human subject during locomotion. We show how a conventional PEEK-cranioplasty provides enough acoustic transmission to perform detailed fUSi of the sensorimotor cortex of the lip, consistently over multiple repetitions, spread out over a period of close to two years. We first describe a personalized 3D-printed fUSi-helmets to fixate the ultrasound probe on the subject’s head, to ensure stability of the probe during walking tasks as well as enable reproducibility of the same 2D-imaging plane across measurements.

Next, we set up an optical tracking pipeline of the subject’s face and ultrasound probe, which allowed for CT/(f)MRI-coregistration of our fUSi-data, as well as tracking of facial movements during functional tasks. Functional tasks focus on sensorimotor activation of the lips and included lip licking, lip pouting and sensory stimulation of the lips. An extensive set of measurements, including several task variations and functional controls, are performed over a period of nearly two years. Our work shows reproducible and consistent fUSi-signal over a period >20 months, with robust acquisitions of functional brain signal even during high-motion scenarios such as walking as shown in this paper. These results demonstrate the feasibility of fUSi for monitoring human brain activity in ecological contexts and serves as an important milestone towards fUSi-based discoveries of the human brain in clinical and neuroscientific context.

## Results

### Subject Recruitment

We recruited two male subjects in their 30s with a PEEK-cranioplasty to participate in our study. Subject #1 received a left-sided hemicraniectomy and PEEK-implant after high-velocity trauma, causing multiple cerebral contusions and post-operative neurological deficits. Most pronounced was the subject’s aphasia, which was considered severe based on baseline linguistic assessment (**Supplementary Data 1**). In addition, although the subject retained the ability for independent walking, he had motor deficits in the right arm and leg, resulting in a right-sided limp. Subject #2 received a PEEK-implant over the right-sided frontotemporal region after surgical removal of a low grade astrocytoma in the right insular region. At time of inclusion, the subject had been tumor progression-free for multiple years. No cognitive, motor or language deficits were reported or objectified at baseline.

Experiments were conducted over a period of nearly 2 years. Subject #1 participated in a total of seven measurements during this period. Subject #2 deceased during the course of the study due to tumor regrowth and participated in a total of two measurement sessions. The results of these measurements will be shown in this manuscript. However, the main portion of the presented data, including the walking experiments, will involve subject #1 only. More details on subject characteristics can be found in **Supplementary Data 1.**

### fUSi-helmet design

To facilitate the reproducibility of our measurements over time, we first designed and tested personalized 3D-printed PLA-helmet to fixate the fUSi-probe with respect to each subject’s brain anatomy. The helmet was based on the subject’s head contour as extracted from MRI-scans and contained two optical geometries necessary for optical tracking (**Figure 1A**). To ensure stability of the probe during functional tasks as well as enable reproducibility of the same 2D-imaging plane across measurements, probe inserts were designed based on targeted brain regions of interests (ROIs) for functional tasks (**Figure 1A**). More details on the pipeline to design and 3D-print personalized helmets and probe inserts can be found in **Supplementary Data 2**.

**Figure 1.**
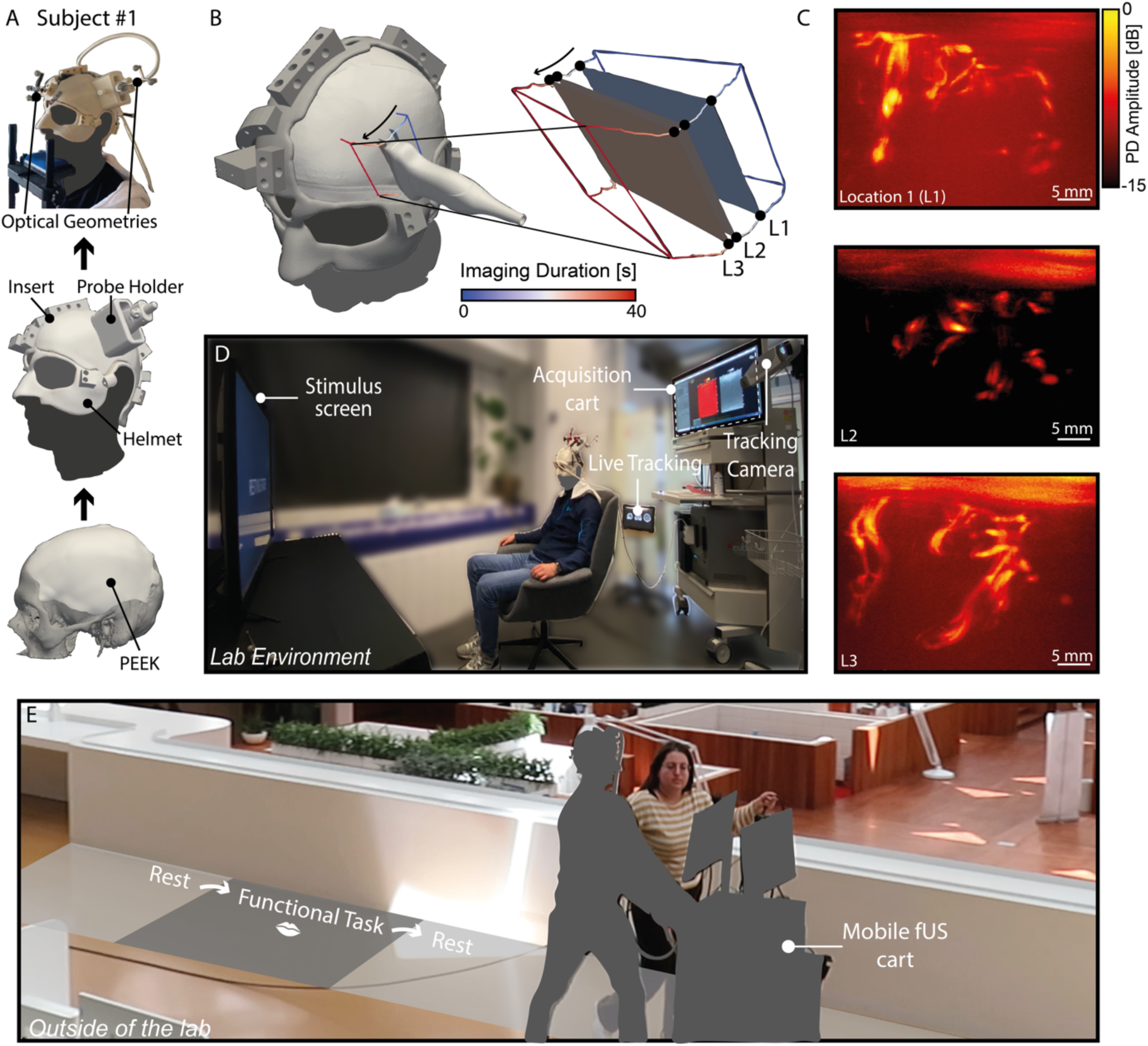
Key components of our experimental set-up. The person depicted in panel E is one of the authors, consenting to their image being displayed in this way.

### Optical tracking and Experimental set-up

The position and orientation of the ultrasound probe (GEL9-D, General Electric, USA) and helmet were tracked continuously using an optical tracking camera (Polaris Vega, Northern Digital Inc., Canada) combined with our custom software (**Supplementary Data 3**). The tracking data was also saved for later use, facilitating offline 3D reconstruction of the ultrasound path, as demonstrated in **Figure 1B**.

fUSi-acquisitions were first performed in our lab-environment dedicated to in-human imaging (**Figure 1C**). Our aim was to create an ecological environment in which subjects could move within the restraints of being tethered by the cord of the ultrasound probe. Once the signal and helmet proved robust, we moved outside of the lab on several occasions using a mobile cart version of our experimental research system for experimental acquisitions during locomotion (**Figure 1D**). During our acquisitions, several data-streams were acquired and stored synchronously, as can be appreciated in **Supplementary Data 4**. These data streams included video recordings of the subject’s face, the movement of which were tracked for functional analyses.

### Validation of Anatomical Localization of the Probe

In both subjects, we designed inserts which positioned the probe and fUSi plane directly over the precentral and postcentral gyrus of the left (in subject #1) (**Figure 2A**) and right (in subject #2) (**Figure 3A**) hemisphere. The precentral and postcentral gyri correspond to the primary motor and sensory cortex respectively, and care was taken to plan the insert over the lower section of the homunculus^34^, traditionally involved in sensorimotor control of the mouth, lip and tongue. Literature contains multiple examples of studies demonstrating the somatotopy of sensorimotor activation of the lips^34,35^, albeit no strong evidence exists for somatotopy of the upper vs. lower lip, or the right vs. left side of the lip, with often heterogenous bihemispheric activation^36–42^. This sensorimotor area was of particular interest, as the anatomical localization of the PEEK in both subjects allowed for easy access. Additionally, we wished to perform a similar task in fUSi as in fMRI for validation purposes, for which the sensorimotor cortex of the mouth seemed particularly useful. Finally, our previous fUSi work in the context of awake neurosurgical procedures involved many iterations of lip pouting or lip licking tasks, which ensured our familiarity with the functional paradigm in the context of fUSi^18,22^. The correct positioning of the fUSi-probe was verified in real-time with our optical tracking system (**Supplementary Data 4**). Co-registration of the Power Doppler images (PDIs) and corresponding MRI-slice confirmed correct localization based on vascular patterns following gyri and sulci (**Figure 2B and 3B**).

**Figure 2.**
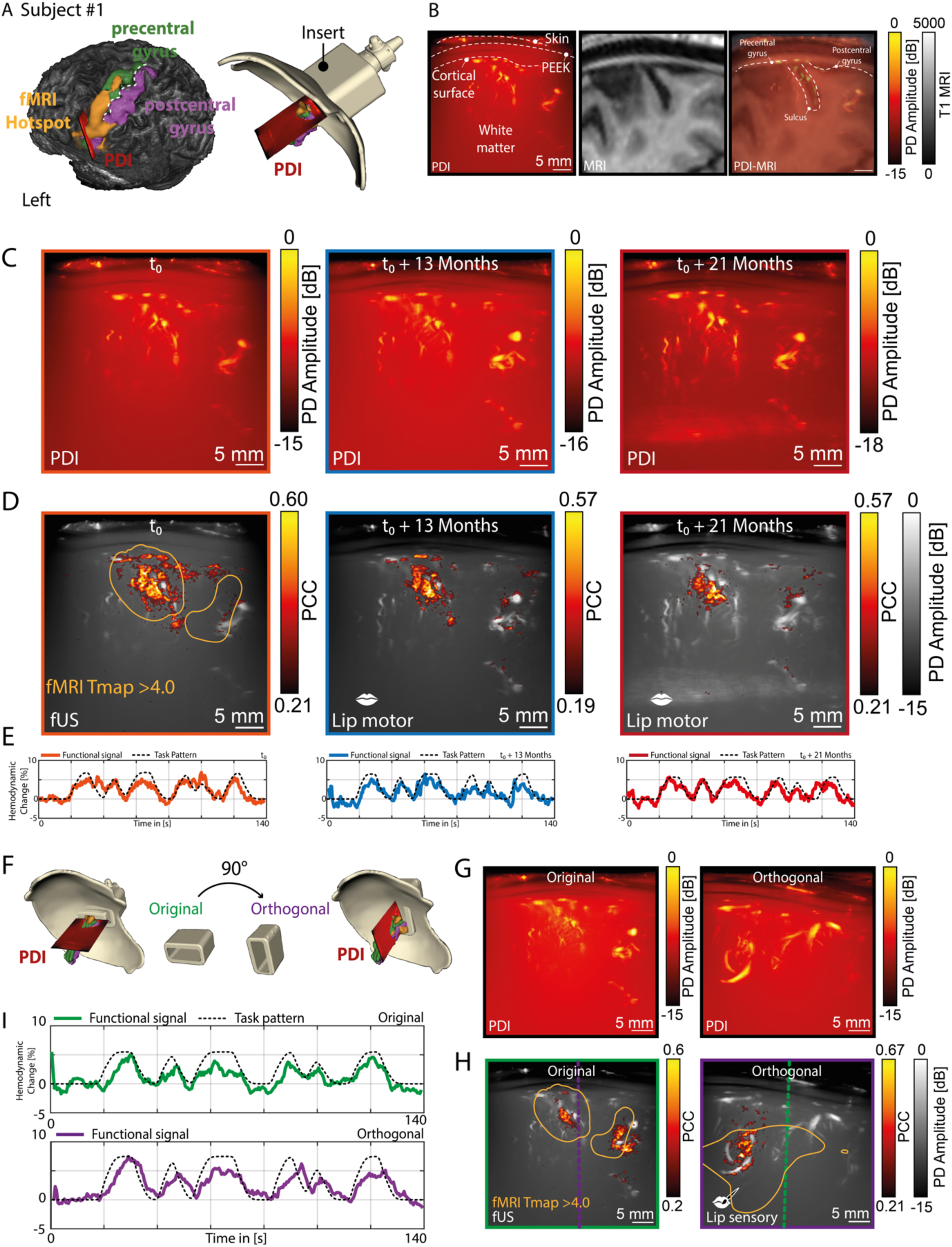
Functional localization and reproducibility of the sensorimotor signal of the lip over time in subject #1. A) The ROI was planned around the sensorimotor cortex of the mouth, overlapping with the fMRI hotspot found during a prior lip pouting task in the same subject. B) The accuracy of the PDI localization could be confirmed both anatomically and by the overlap with the gyri and sulci contour in MRI. C) In subject #1, we were able to revisit the same PDI-plane at three time-points (t_0_, t_0+13 months_, t_0+21 months_), demonstrating the reliability of the personalized helmet and insert combination over time. D) At those same three time-points (t_0_, t_0+13 months_, t_0+21 months_), we were also able to consistently map out functional brain activity during a motor lip pouting task, demonstrating the robustness and reproducibility of the fUSi-signal. The first panel also demonstrates how the functional region as found with fUSi overlapped with the fMRI hotspot as found during a similar lip pouting task (orange contour). E) The corresponding functional signals as found at each of the time points depicted in C and D (t_0_, t_0+13 months_, t_0+21 months_). G) To further study the robustness of the signal, we designed a new orthogonal insert with a probe positioned at a 90-degree angle relative to the original ROI for a lip sensory task (lip brushing). G) The PDI plane of the original vs. the orthogonal insert. H) Functional maps of the original vs the orthogonal insert, with the orange contour showing the relative overlap with the lip pouting task hotspot in fMRI (see also panel A). The green and purple dotted lines in each of the functional maps visualize the intersectional plane with between the original and orthogonal PDI. I) Functional signals corresponding to the functional maps displayed in panel H.

**Figure 3.**
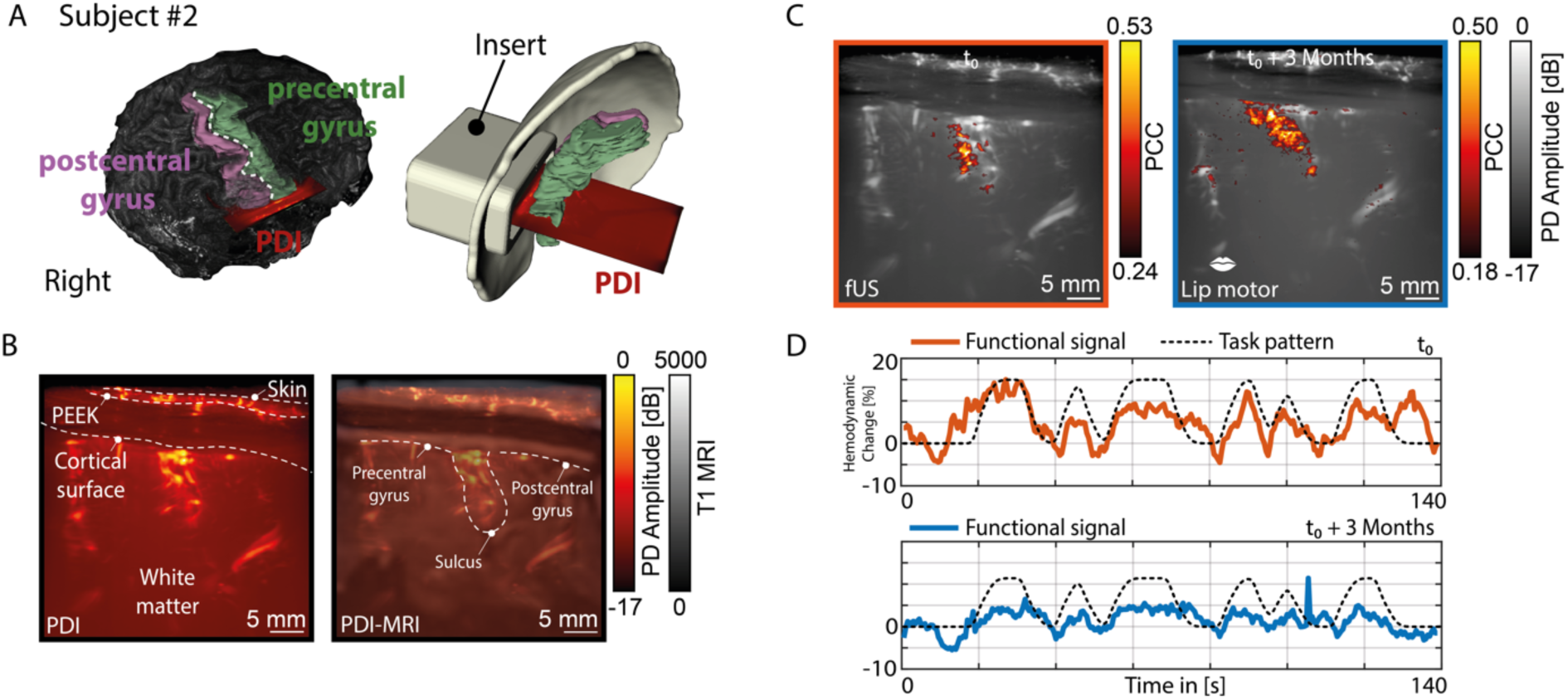
Functional localization and reproducibility of the sensorimotor signal of the lip over time in subject #2. A) The ROI was planned around the sensorimotor cortex of the mouth. B) The accuracy of the PDI localization could be confirmed both anatomically and by the overlap with the gyri and sulci contour in MRI. C) In subject #2, we were able to revisit the same PDI-plane at 2 time-points (t_0_, t_0+3 months_), demonstrating the reliability of the personalized helmet and insert combination over time. At those same two time-points, we were also able to consistently map out functional brain activity during a motor lip pouting task. D) The corresponding functional signals as found at each of the time points depicted in C (t_0_, t_0+3 months_).

The acoustic properties of PEEK are distinctly different than that of brain tissue, leading to distortion of our ultrasound signal through the PEEK-material. In **Supplementary Data 5** we study the nature of these distortions and provide a quick fix and an exact algorithmic solution to obtain undistorted ultrasound images through PEEK so that they can be matched with other modalities such as in our case with fMRI. For the rest of the study where we only study the fUSi data without reference to other modalities we used the quick fix solution which entails image reconstruction using a higher global sound speed to account for the increased sound speed in PEEK with respect to brain tissue.

### Validation of Functional Localization of the Probe using fMRI

Functional mapping using fUSi is accomplished by acquiring a series of sequential power Doppler images (PDIs) to measure changes in cerebral hemodynamics. The spatiotemporal changes between these images serve as a proxy for neural activity through the process of neurovascular coupling^14,19,20^.

We wanted to first confirm the presence of functional activity within our fUS-image prior to expanding our experiments. Therefore, we performed a simple ON-OFF lip pouting (motor) and lip brushing (sensory) task (**Figure 2C-E** and **Figure 3C-D**) in both subjects, to create fUSi-maps depicting functional pixels (*defined as pixels which showed a Pearson Correlation Coefficient (PCC)-value of >3x the standard deviation of the mean PCC value of voxels in a pre-defined noise region*). To perform our PCC-analyses, the stimulus pattern was first convolved with an estimated Hemodynamic Response Function (HRF)^43–47^. As far as we know, there is no data available on the human HRF for fUSi. We therefore estimated an HRF ourselves using 4 training datasets in which the subject performed a lip licking task while walking, similar to what will be shown in **Figure 5**. This training set was obtained one week prior that of the set used for **Figure 5**. The HRF was found the by minimizing the error between the measured fUSi signal and the task time course convolved with the HRF kernel. The HRF kernel itself was modeled as a weighted sum of basis-functions. Specific details of this procedure can be found in **Supplementary Data 6.**

We were able to demonstrate overlap in fUSi-based functional regions as compared to co-registered fMRI-hotspots as found during a prior lip pouting task in fMRI (**Figure 2A,D**). Previous work by our team showed similar consistent overlap between fUSi and fMRI regions in subjects imaged during awake neurosurgical procedures^22^. More details on the functional tasks used for fUSi and fMRI can be found in **Supplementary Data 7.**

Given the limited field of view of our single 2D-plane, we chose to also confirm the robustness of the functional hotspot by designing a new insert for subject #1 with the probe positioned at a 90-degree angle relative to the original ROI, using our optical tracking data. The orthogonally positioned PDI intersected the fMRI hotspot as well and again confirmed the presence of a functional region during a sensory lip brushing task (**Figure 2F-G**).

### Reproducibility and Consistency of the fUSi-signal over time

Both in subject #1 and subject #2, we could reproducibly map functional brain activity during both sensory (lip brushing) and a motor lip pouting task over a period up until 21 months (**Figure 2C-E**, **Figure 3C-D**). This demonstrates how the fUSi-signal could be acquired reproducibly and consistently over longer periods of time using our helmet and insert combination.

### Functional Specificity of the fUSi-signal

The functional region delineated during the motor task (lip pouting, **Figure 2B-E**) localizes primarily in the precentral gyrus, as to be expected. However, the sensory task performed in subject #1 (**Figure 2G-H**, left panel), localizes in both the precentral and postcentral gyrus, indicating potential involvement of both sensory and motor cortex. Although this may raise the question to what extent the hemodynamics-based fUSi-mapping is spatially selective, it should be noted that the primary sensory prominently projects to motor cortex, often eliciting co-activation upon electrical stimulation^48–51^.

To further study the specificity of the functional signal using fUSi, we introduced several task variations in subject #1, as can be seen in **Figure 4**. A classic ON (*lip brushing*) - OFF (*rest*) sensory lip task evoked a reproducible functional fUSi-map with two distinct functional regions (ROI 1 and 2, **Figure 4A**) as was also seen in **Figure 2**. Alternating the ON-task with a different OFF-tasks such as brushing the forehead (**Figure 4B**), the right ear (**Figure 4C**) or the hand (**Figure 4D**) did not significantly alter the functional map.

**Figure 4.**
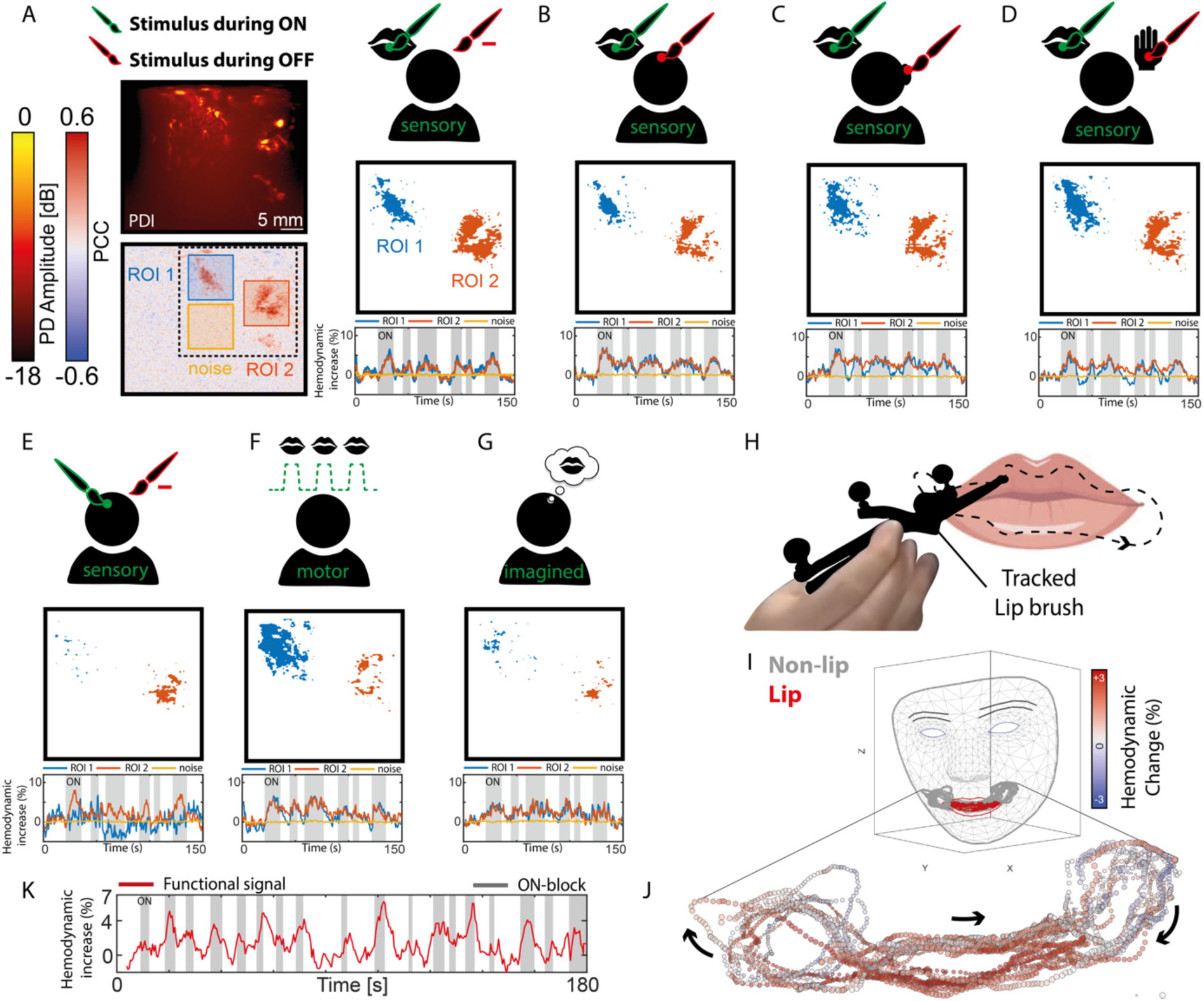
Demonstrating the functional specificity of the fUSi-signal in subject#1. A) A series of task variations in the lip sensory task. A classic ON (lip brushing) - OFF (rest) sensory lip task evokes a reproducible functional fUSi-map with two distinct functional regions (ROI 1 (corresponding to the precental gyrus) and ROI 2 (corresponding to the postcentral gyrus)). The adjacent noise region used for functional thresholding is marked in yellow. The thresholded functional map (showing PCC-values of >3x std of the noise signal only) highlights the two distinct functional regions. B) Alternating the ON-condition (lip brushing) with OFF-task of brushing the forehead, produces a similar functional map. C) Alternating the ON-condition (lip brushing) with OFF-task of brushing the right ear, produces a similar functional map. D) Alternating the ON-condition (lip brushing) with OFF-task of brushing the hand, produces a similar functional map. E) Brushing the forehead only (ON), with a rest-condition in OFF times, leads to far less functional activation, only now present in ROI 2. F) A motor task (lip pouting) in the same task-pattern as shown in panel A does give functional activity, but different in spatial pattern, with are more prominent localization in the precentral gyrus (ROI 1). G) The imagined version of the task shown in panel A resulted in a similar functional map as seen in panels A-D, although less prominent. H) To further study the robustness of our signal, we moved on to a continuous lip brushing task, using optical tracking to determine brush position at each time-point I) Using the MediaPipe library by Google^52^, we could then extract the facial parameters such as the Blendshape coefficients, enabling us to project the lip brushing trace over a mesh representation of the subject’s facial anatomy. J) Plotting the average percentual hemodynamic change in our ROIs as a function of brush location, demonstrates a specific rise in hemodynamic signal only when the lip itself was brushed. K) Functional signal as found in the ROI when the brush was in the lip region (defined as the ‘ON-blocks’).

As a control, we performed a forehead-brushing task only, with no involvement of the lip (**Figure 4E**), where we see close to no activation, with only a small number of functional pixels in ROI2. These findings suggest that the region of the sensorimotor cortex we are imaging seems indeed lip-specific, not involving functional activity of adjacent regions on the sensorimotor homunculus.

A motor task (lip pouting) in the same task-pattern as shown in **Figure 2** did give functional activity in the same region of our ROIs, but different in spatial pattern (**Figure 4F**), with more involvement of the precentral gyrus, as is to be expected. Asking the subject to imagine lip pouting in the ON-times, similar to the imagined task variations performed in fMRI studies^11–13^ (**Figure 4G**), resulted in a similar functional map as seen in panels A-D, although less prominent. Details on the exact specification of these functional tasks can be found in **Supplementary Data 7.**

### Mapping Functional Signal in a Continuous Task

Once the specificity of the functional signal was confirmed, we set out to perform a functional task without a predictable ON-OFF pattern, to further study the robustness of the functional signal. The task consisted of continuous lip brushing by one of the researchers (**Figure 4H**). The tracked brush was moved continuously for several minutes in an unpredictable pattern over the upper and lower lip, as well as adjacent parts of the face. Using the optical tracking of the helmet, as well as the brush itself, we could determine the position of the brush relative to the subject’s facial anatomy accurately at each timepoint of the experiment. Using the MediaPipe library by Google^52^, we could then extract the facial parameters such as the FaceMesh, Blendshape coefficients, and Rotationmatrix from a video file of the face taken during the experiments (more explanation on this in the Methods). As such, we could project the brush trace over a mesh of the subject’s face, allowing us to discern timepoints where the brush touched the lip vs. the adjacent cheeks (**Figure 4I**). Plotting the average percentual hemodynamic change in our ROIs as a function of brush location, demonstrates a specific rise in hemodynamic signal only when the lip itself was brushed vs. the surrounding face (**Figure 4J-K**), again conforming the specificity of the signal.

### fUSi using a mobile cart to facilitate walking

After studying the fUSi-signal extensively in sedentary position using the helmet, we proceeded to an experimental set-up during walking. The ultrasound acquisition system could be pushed by subject #1 while he was still tethered to the acquisition system with the ultrasound probe (**Figure 1E**). The cart contained two separate screens to display the functional task video, as well as the real-time PDIs during acquisition. A long (100m) extension cord was used to allow for a large range of motion. Further details on the acquisition system can be found in the Methods.

During recordings of a little over 1 minute each, the subject (**Figure 5A**) was asked to walk a continuous straight line of a maximum of 30 meters, while pushing the mobile fUSi-cart and following a functional task video displayed on the screen. The subject would walk this continuous straight line in both directions, both away and towards the starting point, to avoid unnecessary delays in successive recording iterations. A camera recorded the subject’s face continuously to allow for post-hoc functional analyses of facial and especially lip movement (**Figure 5B**). The subject was first asked to perform a simple ON-OFF lip licking task while walking, both based on a visual ON-OFF cue in video-form, as well as an audio ON-OFF cue, given verbally by the experimenter based on certain landmarks on the ground (**Figure 5A**). More details on the functional tasks used can be found in **Supplementary Data 7**.

**Figure 5.**
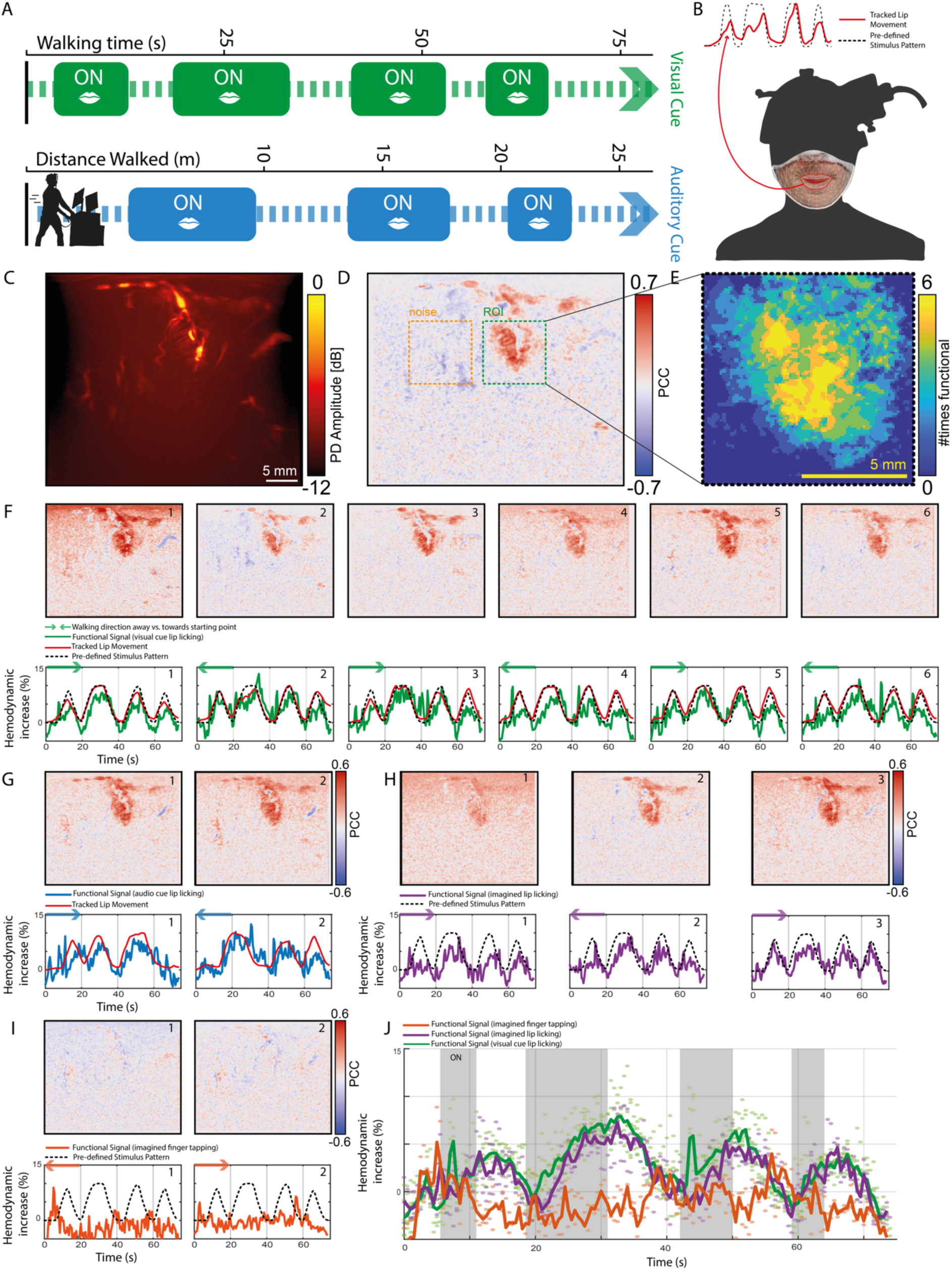
fUSi-signal acquired during walking. A) Overview of the experimental conditions, where ON-blocks (lip licking, finger tapping or imagined lip licking) were either introduced with a visual cue (task video shown on screen) or audio cue (ON and OFF indicated verbally by experimenter based on landmarks on the ground). B) Example video footage taken from the subject during walking and used to analyze facial movements post-hoc using the MediaPipe library by Google^52^. The red trace represents an example trace of the tracked lip movements which could be extracted from the video material. C) Example of the average PDI as acquired during the walking imaging sessions. D) Example of a fUSi-map acquired during a visual lip licking task (see panel A), with the ROI marked in green. The adjacent noise region used for functional thresholding is marked in yellow. E) A total of six iterations of the task shown in panel D were performed. Here we summarize the presence of functional pixels (defined as >4x std of the noise signal) in the ROI across these 6 iterations. Pixels with a higher summarized value were present in a higher number of iterations of this particular task, highlighting the robustness of their functional involvement. F) Six example iterations of the same functional task as shown in panel D (visual based lip licking), showing the reproducibility and robustness of the functional signal captured during locomotion. The arrow directions on the function signal plots signify the walking direction (either towards or away from the starting point) during the measurements. G) Example traces of the auditory-based lip licking task. H) Functional maps and functional signal in three iterations of the imagined lip licking task show similar activation as in panel E, although slightly less pronounced in signal amplitude. I) As a control condition, imagined finger tapping in the same task pattern did not evoke any functional signal. I) Summarizing scattering plot showing the functional signal in the lip licking task (green), as well as in the imagined lip licking task (purple), albeit the latter being lower in signal amplitude. The summarizing scatter plot confirms the lack of functional signal in the imagined finger tapping task (orange), confirming the specificity of the signal.

### Reproducible and Consistent fUSi-Signal during Walking

We were able to produce high-quality PDIs (**Figure 5C**) and functional maps (**Figure 5D-E**) while acquiring a lip licking functional signal during locomotion. These functional maps were consistent and reproducible over multiple iterations, both when using a visual cue (**Figure 5F**), as well as an audio cue (**Figure 5G**). Close to no motion correction was necessary during the preprocessing of our PDI-stacks, indicating a high level of in-plane stability during image acquisitions (**Supplementary Data 8**). Interestingly, a similar signal could be acquired when asking the subject to imagine lip licking while looking at the functional video, *without* actively performing the task (**Figure 5H**). A control task with imagined finger tapping, did not result in any functional activation, confirming again the specificity to the imagined lip pouting signal (**Figure 5I**). A summarizing scatter plot in **Figure 5J** demonstrates the specificity of the lip-related functional signal acquired during walking.

## Discussion

This paper demonstrates robust and reproducible fUSi of human brain activity during walking. Using a conventional ultrasound probe and a 3D-printed personalized helmet, we were able to consistently revisit the same ROIs in two subjects, despite the challenges that come with repeated 2D-imaging. We were able to demonstrate activation of the sensorimotor cortex of the mouth with both motor and sensory tasks, confirmed by co-localization with the respective fMRI hotspot. The functional maps and underlying functional signals were reproducible over a period of up to 21 months. We performed multiple variations of the sensory lip task, confirming the functional region’s lip-specific responsiveness. We were also able to acquire robust and reproducible functional data outside of the controlled lab environment, with our subjects walking during our acquisitions, without facing motion-related problems.

Compared to the neurosurgical setting our team is accustomed to, having access to tethered but freely moving subjects who can be imaged over multiple sessions, was a game-changer. Many of the control or repetition measurements we show in this manuscript are simply not possible in the intra-operative context given the limited time as well as constraints on the stamina of the awake subject. However, in light of technology development and validation, repetition is key. Each of our functional measurements produced consistent results, even with months between measurement sessions, demonstrating the robustness of our results.

One striking observation during our experiments is the apparent specificity of the lip-related activation of the sensorimotor cortex as imaged by fUSi, both in the active as well as the imagined condition. Although literature tells us we could rightfully expect our hemodynamic technique to spatially discriminate representation sites which are in close proximity in the sensorimotor homunculus^37,53^, our first fUSi observations inspire further research. For example, it would be interesting to further unravel the ‘imagined’ functional task conditions shown in this paper. Although fMRI literature relies heavily on motor imagery paradigms to study sensorimotor brain activity^11–13,54^, it is still striking how similar our ‘imagined’ maps are to the actual motor paradigm, especially in **Figure 5F** and **H**. One explanation could be our subject’s misinterpretation of the task: despite us seeing no detectable movement of the tongue or lip on video recordings during the imagined conditions, the subject could have performed a similar version of the task within the oral cavity and the mouth closed.

Similarly, the consistent co-activation of what would be precentral gyrus during a sensory task as shown in **Figures 2** and **3**, as well as the persisting functional region found when performing sensory stimulation of the forehead only, as shown in **Figure 4E**, warrant further investigation. Similar to the very first homunculus mapped out by Penfield^34^ in the 1930s using electrical stimulation, it would be interesting to pinpoint how much spatial selectivity we can actually achieve when imaging human cortex using fUSi, a hemodynamics-based signal. These observations can be incredibly informative in light of future ambitions such as fUSi-guided decoding of brain signals for Brain Computer Interfaces (BCIs). The first examples in nonhuman primates using fUSi-based decoding of motor intention, seem promising^55,56^.

Before any of the above can actually be accomplished, we need to move away from our 2D-context, which is spatially limited sensitive to out-of-plane movements, towards the use of real-time 3D-probes. To overcome the challenges of 2D-imaging in this study, we used our custom 3D-printed helmet 1) as a reference for the relative probe position to the subject’s neuro-anatomy, 2) as a frame to physically stabilize the probe over an ROI during the task, and 3) as a means to be able to revisit the exact same 2D-plane over multiple imaging sessions. Although the current helmet facilitated all three functions, a 3D-fUSi capable probe is likely to further ameliorate artifact correction due to motion, out of plane movements, and/or inconsistencies over multiple imaging sessions. What is more, 3D-fUSi will allow us to perform more dynamic mobile mapping of the human brain, opening up possibilities such as dynamic decoding of the sensorimotor cortex during locomotion, which could be extremely valuable in monitoring and active guidance of functional plasticity during rehabilitation programmes in tumor or neurotrauma patients. For this application, however, overcoming the challenge of real-time processing of vast amounts of 3D fUSi-data, will be particularly essential.

Ultimately, our ambition would be to image not just through cranioplasties such as PEEK, but transcranially with the same or even better imaging quality as demonstrated in this paper. The current inability to do so is one of the major limitations of ultrasound-based techniques in humans. Although literature shows how this limitation may be overcome by using contrast-agents and performing so called transcranial ultrasound localization microscopy (ULM)^57,58^, ULM in functional settings is still unfavorable given the need for intravenous microbubble delivery and long time windows for data acquisitions.

Nevertheless, our work confirms the compatibility of unmodified clinical-grade cranioplastes with fUSi experiments. Part of the appeal of PEEK in particular, in addition to its sonoluscency, is its already widespread clinical use, which could warrant cranioplasties as a new clinical standard of care for e.g. post-operative monitoring purposes. In the setting of traumatic brain injury, for example, continuous acoustic access to the brain could add an additional imaging modality to monitor functional connectivity in comatose patients in the ICU, aiding in prediction of functional outcome and treatment decisions accordingly. Currently, only expensive and logistically challenging fMRI-scans or EEG-measurements are available in the ICU for these purposes^59,60^.

Apart from just for functional imaging, the structural images we could make through PEEK using a commercially available ultrasound machine (see **Supplementary Data 9**), could open up a new way of post-operative monitoring of tumor (re)growth, without the need to rely on periodical expensive MRI-scans. An avenue which so far seems to have remained mostly undiscovered in the clinical context^29^.

Our current paper follows shortly after the study of Rabut et al.^33^ published in *Science Transtlational Medicine*^33^, which was developed in the same time-frame. In their paper titled ‘*Functional ultrasound imaging of human brain activity through an acoustically transparent cranial window*’, the authors demonstrate their development of a thinned, experimental PMMA-cranioplasty, implanted in a subject with a hemicraniectomy after trauma. The authors show their ability to capture functional activity during a gaming and guitar strumming task. What sets our study apart, is the fact that we demonstrate in-human mobile fUSi: we can capture functional brain activity in a walking human. Furthermore, the longer timespan of our measurements has allowed for an in-depth analysis of the functional validity of the fUSi-signal and includes many repetitions of datasets and functional task verifications to ensure the robustness, reliability and the reproducibility of the signal. The different task variations we designed throughout the years, allow us to confidently couple our functional signal to meaningful brain activity relating to sensorimotor control of the mouth. The swift follow-up of the Rabut et al.^31^ and our paper does send out one clear, unified message: using fUSi through already available, clinical grade cranioplasties provides unique access to human brain functionality.

In this paper we demonstrate for the first time that fUSi allows us to map human brain functionality during walking in subjects with sonolucent PEEK cranioplasties, at depths of multiple centimeters, in a robust and reproducible fashion. These observations further fuel the field to consider fUSi as more than just an intra-operative tool, pushing towards the development of mobile, in-human fUSi as a new means to unravelling the human brain in clinical and neuroscientific context.

## Methods

### Subject Recruitment

Two subjecs with a PEEK-cranioplasty were recruited from the Department of Neurosurgery of the Erasmus MC in Rotterdam. Prior to inclusion, written informed consent was obtained in line with the National Medical-Ethical Regulations (MEC-2019-0689 and MEC-2022-0087, NL80307.078.22).

### Baseline Linguistic and Cognitive Assessment

To determine baseline linguistic and cognitive abilities op both subjects, a standard clinical test battery including language tests such as the DIMA and DuLIP (Diagnostic Instrument for Mild Aphasia and Dutch Linguistic Intraoperative Protocol) and cognitive tests such as the Trailmaking-test were performed by trained clinical linguist (*DS*) (**Supplementary Data 1**).

### 3D-printed Personalized Helmet

We designed a personalized 3D-printed PLA-helmet to fixate the fUSi-probe with respect to the subject’s brain anatomy. The helmet was based on the subject’s head contour as extracted from MRI-scans and contained two optical geometries necessary for optical tracking (Northern Digital Inc., Canada). A step-by-step explanation of the production pipeline can be found in **Supplementary Data 2**.

### Optical Tracking and Ultrasound Path Reconstruction

Position and orientation of the fUSi-probe and helmet was tracked continuously using an NDI Polaris Vega optical tracking system (SN P9-04539, Northern Digital Inc., Canada), which was configured to track infra-red reflective reference geometries attached to the fUSi-probe and helmet. Custom software was designed to record the tracking information featuring six degrees of freedom (DOF) at an average rate of 20 Hz.

To be able to visualize the fUSi-probe’s position relative to the subject’s brain in real-time during our experiments, we utilized the tracking information in conjunction with custombuild software using the Visualization Toolkit (VTK)^61^. The tracking data was also saved for later use, facilitating offline 3D reconstruction of the ultrasound path, as demonstrated in **Figure 1B**. More details on our optical tracking pipeline can be found in **Supplementary Data 3**.

### Experimental Context

fUSi-acquisitions were mostly performed in our lab-environment dedicated to in-human imaging (**Figure 1C**). Our aim was to create an ecological environment in which subjects could freely move within the restraints of being tethered through cord of the ultrasound probe. On several occasions, we moved our mobile experimental research system to an external location (the hallway of our lab building) for experimental acquisitions during locomotion (**Figure 1D**). During our acquisitions, several data-streams (ultrasound, optical tracking, video) were acquired and stored synchronously, as can be appreciated in **Supplementary Data 4**.

### fUSi-acquisitions

fUSi-acquisitions were performed using an experimental research system (Vantage-256, Verasonics) interfaced with a 9L-D linear array transducer (GE, 5.3 MHz). For all scans we acquired continuous angled plane wave acquisition (10 equally spaced between −12 and 12 degrees) with a PRF of 800 Hz. The average ensemble size (number of frames used to compute one Power Doppler Image (PDI)) was set at 200 frames from which the live PDIs were computed, providing a live Doppler FR ranging of 4 Hz. The PDIs as well as the raw, angle compounded beamformed frames were stored to a fast PCIe SSD hard disk for offline processing purposes.

### fUSi Mobile Cart

To facilitate measurements during walking, we loaded the Vantage-256 and analysis PC onto a small cart, which could be pushed by the subject himself. Electrical power was provided through a long 100 meter long extension cord and a safety isolating transformer. The cart contained two screens, one facing the subject and showing the functional video, and one facing the experimental team showing real-time PDIs during acquisition. The mobile cart was equipped with several video cameras, which were captured synchronously with the PDI-data (see **Supplementary Data 4**) using OBS Studio screen recorder (OBS Studio Contributors).

### (f)MRI-acquisitions

Each subject underwent a single fMRI-scan of max. 60 minutes prior to the fUS acquisitions. MR imaging was performed at 3.0T with an 8-channel head coil (Discovery MR750, GE Healthcare, Milwaukee, WI, US). Whole brain functional MR images were obtained with a single shot T2* weighted EPI sequence sensitive to blood oxygenation level dependent (BOLD) contrast with the following parameters: repetition time (TR) = 3000 ms, echo time (TE) = 30 ms, flip angle = 90°, acquisition matrix = 96 x 64, field of view (FOV) = 240 x 180 mm^2^. We acquired 54 slices with a slice thickness of 2.2 mm and 0.3 mm gap. Additionally, one higher temporal resolution scan was performed with TR = 1500 ms and 27 slices covering the brain parenchyma underlying the skull bone defect. All functional data acquisition started with 5 dummy scans, which were discarded from further analysis. Subjects received instructions and practiced the fMRI task together with a researcher prior to MRI scanning. During scanning, stimuli were visually presented outside the scanner onto an MRI compatible monitor that was visible with a mirror mounted on the head coil. Additionally, a high-resolution 3D T1-weighted inversion recovery fast spin gradient recalled echo (IR FSPGR) structural MRI was acquired in the axial plane with the following parameters: TR = 7.93 ms, TE = 3.07 ms, inversion time = 450 ms, flip angle = 12°, acquisition matrix = 240 x 240, FOV = 240 x 240 mm^2^. 176 contiguous slices were acquired with a slice thickness of 1 mm.

### Functional Paradigms

Based on the anatomical localization of the PEEK in both subjects, as well as the wish to perform a similar task in fMRI for validation purposes, we chose to focus on the sensorimotor cortex of the mouth using motor (lip pouting) and sensory (lip brushing) tasks. For the motor lip pouting task, the subject was asked to follow a task video in which lip pouting was demonstrated during the ON-blocks. For the sensory task, one of the researchers (SS) stimulated the subject’s lips using an optically tracked, fine-haired brush during ON-times. In the majority of the sedentary, fUSi-based functional tasks displayed in this manuscript, a 140 second task-pattern was used with randomized ON-OFF blocks ranging between 4.1 – 15.4 seconds. For the fMRI-acquisitions, longer ON-OFF blocks of 30 seconds each were used, in parallel with our in-house clinical fMRI acquisition schemes. The functional task used during walking focused on lip licking specifically, and involved two task variations, with a total task duration of 74 seconds each. Detailed descriptions of the functional task patterns used during the fUSi- and fMRI-acquisitions can be found in **Supplementary Data 7**.

### fMRI-data processing

fMRI-analysis was performed offline using Statistical Parametric Mapping (SPM8, Functional Imaging Laboratory, UCL, UK) implemented in Matlab (vR2015b). For each subject, we first spatially realigned all fMRI-images and co-registered these images to the individual’s T1-weighted image, using a rigid body transformation as implemented within SPM8. Functional images were smoothed with a 3D Gaussian Full Width Half Maximum (FWHM) filter of 6 × 6 × 6 mm^3^. All fMRI data were analyzed using the general linear model (GLM), by modelling the experimental and the control conditions in a blocked design (see **Supplementary Data 7** for exact task patterns). The blocks were convolved with the hemodynamic response function (HRF), corrected for temporal autocorrelation and filtered with a high-pass filter of 128 s cut-off. Motion parameters were included in the model as regressors of no interest to reduce potential confounding effects of motion. Individual t-contrast images for the experimental versus control condition were generated and thresholded individually at approximately 60% of the maximum t-value. Resulting thresholded images were projected on the 3D T1-weighted image and visually checked by a neuroradiologist with >20 years’ experience (MS) with fMRI to assess that expected activation patterns were detected, with threshold adaptation if necessary.

### Estimating the fUSi Hemodynamic Response Function (HRF)

We estimated a fUSi-specific HRF on the basis of 4 training datasets in which the subject performed a lip licking task while walking, similar to what is shown in **Figure 5**. This training set was obtained one week prior that of the set used for **Figure 5**. The HRF was found the by minimizing the error between the measured fUSi signal and the task time course convolved with the HRF kernel. The HRF kernel itself was modeled as a weighted sum of basis-functions. Specific details of this procedure can be found in **Supplementary Data 6**.

### Extracting Lip Movements from Video using Blendshape

All video streams were synchronously recorded at 60 frames-per-second using the open-source Broadcast Software named OBS Studio (https://obsproject.com/). In the postprocessing stage we separated the individual streams again into various sub-videos (face, cue, etc.) using a Matlab script, with FFMPEG for lossless splitting. We then used the MediaPipe library by Google^52^ to extract the facial parameters such as the FaceMesh, Blendshape coefficients, and Rotationmatrix from a video file of the face. These parameters were obtained using a Python program and were saved in an HDF5 file to facilitate further processing in conjunction with the overall fUSi processing in Matlab. We used the “mouthSmileRight” and/or “mouthSmileLeft” BlendShape coefficient for the generation of the stimulus signal. Prior to correlation we first used a moving median filter of .5 second (31 frames) to remove outliers and fill in missing datapoints after which we convolved the output with our estimated hemodynamic response function. Careful inspection between the video and the BlendShape coefficient revealed a 0.4 delay of the coefficient lagging the video. We accounted for this delay in the further processing and plotting.

### Ultrasound Distortion Correction

We are able to visualize functional activity through the PEEK-cranioplasty, however, it is necessary to increase the bulk sound speed used in the delay-and-sum reconstruction to compensate for the higher sound speed of the implant layer. This results in a warping of the reconstructed vasculature and can generate errors when co-registering to structural MRI. We tested whether it is possible to correct this warping by segmenting the skull implant from the reconstructed b-mode images and using a simple multi-layer ray-tracing model to update the delays for each voxel. We found that this approach resulted in better alignment to MRI, as demonstrated in **Supplementary Data 5**.

### fUSi-data processing

All the Power Doppler Images (PDIs) and associated results are obtained using post processing of the continuous ultrafast ultrasound data that was recorded to disc during the experiments. Although the variety in experiments and datasets would benefit from tailored processing we chose, for the sake of clarity, to have the same processing pipeline and parameters for all datasets shown, except for **Figure 4J** where additional conditioning of the signal was necessary to obtain the desired result. Every PDI was computed using an ensemble of 800 ultrafast ultrasound frames, with a half overlap between consecutive frames yielding a framerate of 2 Hz. The Doppler signal was obtained using an SVD rank reduction technique that works by removing the first dominant singular vectors which span the stationary tissue signal, from the ensemble of frames, leaving only the blood signal. For all sets we removed the first 56 vectors (7%) and the last 40 vectors (5%) to reduce noise. After this Doppler filtering step, we discarded 40 Doppler frames with the highest median absolute deviation (MAD) per frame. The remaining 760 frames where subsequently interpolated onto a 100 µm grid using zero-padding in the frequency domain. The resulting PDI was then computed by averaging the magnitude for every complex pixel signal over all remaining frames. We then checked on outlier PDIs by computing the MAD of every PDI with respect to the median over all PDIs. PDIs with a MAD score of 2 x 1.48 were replaced with a median PDI^62^. This procedure only affected a few datasets that contained substantial motion. Subsequent sub-pixel motion compensation for every PDI with respect to the median PDI was applied using a cross-correlation technique that can be efficiently computed in the frequency domain^63^. The standard deviation of the motion in the x and z direction for all datasets shown in **Figure 5** are respectively 60 and 10 micrometers. Detailed plots of the motion for these sets are shown in **Supplementary Data 8**. Before computing the PDI by averaging per pixel the absolute squared signal over time we performed up-sampling of the ultrasound frames in the frequency domain to a grid-spacing of 100 micrometer in both the depth(z) and width(x) dimension^18^.

For every pixel in the PDI, the Pearson Correlation Coefficient (PCC) ‘r’ was computed between the stimulus signal (convolved with our estimated fUSi HRF, see above) and pixel intensity over time. A noise region was defined in each PDI, where no vascular or response signal was to be expected. Functional pixels were identified as those with PCC-values > 3 x std of the PCC values in the noise region. We produced ‘overlay-figures’, where functional pixels were displayed over the mean-PDI (*grayscale*).

The average hemodynamic time traces or ‘functional signals’ that are shown in Figures 2 to 5 depict the relative change with respect to the baseline signal which in our case is defined as the mean signal amplitude over the first 10 seconds^62^.

### Continuous lip brushing task

For **Figure 4J** we mapped the hemodynamic signal to the brush position with respect to the subject’s face. For this purpose, we first performed an on-off test similar to what is shown in Figure 4A in order to identify the functionally significant pixels. These pixels were then used in the continuous brushing test. The average hemodynamic time trace was computed as described above and was further smoothed using a 5 second moving median filter in order to make the mapping more robust. An average hemodynamic delay obtained from our estimated hemodynamic response function was added to the tracking data time vector to ensure a hemodynamically meaningful mapping between the tracking data and ultrasound data. The hemodynamic time trace then was interpolated to the same time sampling as the tracking data. For this experiment, where we focus on the lips and a portion of the cheeks, we only used the x and y coordinates of the tracking data as the variation in depth (z-dimension) was very minimal. These three vectors (x and y tracking coordinates and hemodynamic signal) where then used for the scatterplot shown in **Figure 4J** were every x,y coordinate gets a dot which is colored by the amplitude of the hemodynamic signal. Additionally, the size of every dot is scaled by the magnitude of the hemodynamic signal.

## Supplementary Materials

Supplementary Data 1 – Subject Characteristics

Supplementary Data 2 – Helmet Fabrication Pipeline

Supplementary Data 3 – Optical Tracking Pipeline

Supplementary Data 4 – Parallel Datastreams recorded

Supplementary Data 5 – Ultrasound Distortion Correction

Supplementary Data 6 – Hemodynamic response function estimation

Supplementary Data 7 – Overview of functional tasks used during fUSi and fMRI

Supplementary Data 8 – Average motion during walking tasks

Supplementary Data 9 – Conventional ultrasound images through PEEK

Supplementary Data 10 – Datasheet

## Materials & Correspondence

Requests should be addressed to Dr. Pieter Kruizinga, corresponding author.

## Data availability

Data is available upon reasonable request to the authors.

## Acknowledgements

The authors would like to thank Ellen Collée for her involvement in the neurolinguistic screening of subjects and the fMRI-scanning. The authors would like to additionally thank members of the radiology team of the Erasmus MC for facilitating the fMRI measurements. Finally, the authors would like to thank both subjects in this study for their time and efforts. This work was supported was supported by the NWO-Groot grant of The Dutch Organization for Scientific Research (NWO), awarded to CUBE (Center for Ultrasound and Brain-Imaging @ Erasmus MC), by NWO-TTW-OTP (TOUCAN), TKI-LSH (RELAY) & 4DBrain). CIDZ and SS were funded by the Dutch Organization for Medical Sciences (ZonMw), Life Sciences and DBI2 (NWO), ERC-advanced, INTENSE (LSH), and BIG of Erasmus MC.

## Author contributions

SS and PK came up with the study design. SS and AV were involved in the subject recruitment. SS and GS were responsible for the helmet design. SS, LV, FM, BG, BK and PK were responsible for fUSi-data acquisition. SS and MS were responsible for fMRI-data acquisition. DS was responsible for linguistic assessment of the subjects. SS, LV, FM, MB, PK were involved in general data-analysis. BH was responsible for the HRF estimations. MB was responsible for the ultrasound distortion correction. SS, LV, FM and PK were involved in data-interpretation. SS and PK wrote the first draft of the manuscript. All authors provided input for and approved the final draft of the manuscript.

## Notes

### Competing Interest Statement

The authors have declared no competing interest.

## References

1. Stangl, M., Maoz, S. L. & Suthana, N. Mobile cognition: imaging the human brain in the ‘real world’. Nat. Rev. Neurosci. 24, (2023).

2. Aliko, S., Huang, J., Gheorghiu, F., Meliss, S. & Skipper, J. I. A naturalistic neuroimaging database for understanding the brain using ecological stimuli. Sci. Data 7, (2020).

3. Shin, J. H. et al. Wearable EEG electronics for a Brain–AI Closed-Loop System to enhance autonomous machine decision-making. npj Flex. Electron. 6, (2022).

4. Sakai, J. Functional near-infrared spectroscopy reveals brain activity on the move. Proc. Natl. Acad. Sci. U. S. A. 119, (2022).

5. Vidal-Rosas, E. E., von Lühmann, A., Pinti, P. & Cooper, R. J. Wearable, high-density fNIRS and diffuse optical tomography technologies: a perspective. Neurophotonics 10, (2023).

6. DeVore, H. et al. High Performance Wearable Diffuse Optical Tomography with 2x2 Source-Detector Modules. in Bio-Optics: Design and Application in Proceedings Biophotonics Congress: Optics in the Life Sciences 2023, OMA, NTM, BODA, OMP, BRAIN 2023 (2023). doi:10.1364/BODA.2023.JTu4B.21

7. Duraivel, S. et al. High-resolution neural recordings improve the accuracy of speech decoding. Nat. Commun. 14, (2023).

8. Topalovic, U. et al. A wearable platform for closed-loop stimulation and recording of single-neuron and local field potential activity in freely moving humans. Nat. Neurosci. 26, (2023).

9. Fred, A. L. et al. A Brief Introduction to Magnetoencephalography (MEG) and Its Clinical Applications. Brain Sciences 12, (2022).

10. Pasquini, L., Peck, K. K., Jenabi, M. & Holodny, A. Functional MRI in Neuro-Oncology: State of the Art and Future Directions. Radiology 308, (2023).

11. Jahn, K. et al. Brain activation patterns during imagined stance and locomotion in functional magnetic resonance imaging. Neuroimage 22, (2004).

12. Blumen, H. M., Holtzer, R., Brown, L. L., Gazes, Y. & Verghese, J. Behavioral and neural correlates of imagined walking and walking-while-talking in the elderly. Hum. Brain Mapp. 35, (2014).

13. Boyne, P. et al. Functional magnetic resonance brain imaging of imagined walking to study locomotor function after stroke. Clin. Neurophysiol. 132, (2021).

14. Macé, E. et al. Functional ultrasound imaging of the brain. Nat. Methods 8, 662–664 (2011).

15. Deffieux, T., Demene, C., Pernot, M. & Tanter, M. Functional ultrasound neuroimaging: a review of the preclinical and clinical state of the art. Curr. Opin. Neurobiol. 50, 128–135 (2018).

16. Imbault, M., Chauvet, D., Gennisson, J. L., Capelle, L. & Tanter, M. Intraoperative Functional Ultrasound Imaging of Human Brain Activity. Sci Rep 7, 7304 (2017).

17. Soloukey, S. et al. Functional imaging of the exposed brain. Front. Neurosci. 17, (2023).

18. Soloukey, S. et al. Functional Ultrasound (fUS) During Awake Brain Surgery: The Clinical Potential of Intra-Operative Functional and Vascular Brain Mapping. Front. Neurosci. 13, 1384 (2020).

19. Nunez-Elizalde, A. O. et al. Neural correlates of blood flow measured by ultrasound. Neuron 110, 1631–1640.e4 (2022).

20. Iadecola, C. The Neurovascular Unit Coming of Age: A Journey through Neurovascular Coupling in Health and Disease. Neuron 96, 17–42 (2017).

21. Demene, C. et al. Functional ultrasound imaging of brain activity in human newborns. Sci. Transl. Med. 9, (2017).

22. Soloukey, S. et al. Human brain mapping using co-registered fUS, fMRI and ESM during awake brain surgeries: A proof-of-concept study. Neuroimage 283, 120345 (2023).

23. Pinton, G. et al. Attenuation, scattering, and absorption of ultrasound in the skull bone. Med. Phys. 39, (2012).

24. Meggyesy, M. et al. First Experience with Postoperative Transcranial Ultrasound Through Sonolucent Burr Hole Covers in Adult Hydrocephalus Patients. Neurosurgery 92, (2023).

25. Zhang, J. et al. The application of polyetheretherketone (PEEK) implants in cranioplasty. Brain Res. Bull. 153, 143–149 (2019).

26. Spena, G., Guerrini, F., Grimod, G., Salmaggi, A. & Mazzeo, L. A. Polymethyl Methacrylate Cranioplasty Is an Effective Ultrasound Window to Explore Intracranial Structures: Preliminary Experience and Future Perspectives. World Neurosurg. 127, e1013–e1019 (2019).

27. Carlson, J. E., Van Deventer, J., Scolan, A. & Carlander, C. Frequency and temperature dependence of acoustic properties of polymers used in pulse-echo systems. Proc. IEEE Ultrason. Symp. 1, 885–888 (2003).

28. Signorelli, F. et al. Bedside Ultrasound for Ventricular Size Monitoring in Patients with PEEK Cranioplasty: A Preliminary Experience of Technical Feasibility in Neurotrauma Setting. Neurocrit. Care 37, 705–713 (2022).

29. Mursch, K. & Behnke-Mursch, J. Polyether Ether Ketone Cranioplasties Are Permeable to Diagnostic Ultrasound. World Neurosurg. 117, 142–143 (2018).

30. Belzberg, M. et al. Sonolucent Cranial Implants: Cadaveric Study and Clinical Findings Supporting Diagnostic and Therapeutic Transcranioplasty Ultrasound. J Craniofac Surg 30, 1456–1461 (2019).

31. Rossitto, C. P. et al. Transcranioplasty Ultrasonography Through a Sonolucent Prosthesis: A Review of Feasibility, Safety, and Benefits. World Neurosurgery 178, (2023).

32. Williams, A. L. et al. Letter: The Role of Sonolucent Implants in Global Neurosurgery. Neurosurgery 94, (2024).

33. Rabut, C. et al. Functional ultrasound imaging of human brain activity through an acoustically transparent cranial window. Sci. Transl. Med. 16, eadj3143 (2024).

34. Penfield, W. & Boldrey, E. Somatic motor and sensory representation in the cerebral cortex of man as studied by electrical stimulation. Brain 60, 389–443 (1937).

35. Zhao, M., Marino, M., Samogin, J., Swinnen, S. P. & Mantini, D. Hand, foot and lip representations in primary sensorimotor cortex: a high-density electroencephalography study. Sci. Rep. 9, (2019).

36. Muret, D., Root, V., Kieliba, P., Clode, D. & Makin, T. R. Beyond body maps: Information content of specific body parts is distributed across the somatosensory homunculus. Cell Rep. 38, (2022).

37. Grabski, K. et al. Functional MRI assessment of orofacial articulators: Neural correlates of lip, jaw, larynx, and tongue movements. Hum. Brain Mapp. 33, 2306–2321 (2012).

38. Baumgartner, C., Barth, D. S., Levesque, M. F. & Sutherling, W. W. Human hand and lip sensorimotor cortex as studied on electrocorticography. Electroencephalogr. Clin. Neurophysiol. Evoked Potentials 84, (1992).

39. Disbrow, E. A., Hinkley, L. B. N. & Roberts, T. P. L. Ipsilateral Representation of Oral Structures in Human Anterior Parietal Somatosensory Cortex and Integration of Inputs across the Midline. J. Comp. Neurol. 467, (2003).

40. Fox, P. T. et al. Location-probability profiles for the mouth region of human primary motor-sensory cortex: Model and validation. Neuroimage 13, (2001).

41. Kitayama, C. et al. Magnetoencephalographic evaluation of repaired lip sensation in patients with cleft lip. PLoS One 17, (2022).

42. Mogilner, A. et al. Neuromagnetic studies of the lip area of primary somatosensory cortex in humans: evidence for an oscillotopic organization. Exp. Brain Res. 99, (1994).

43. Kotti, S. E., Erol, A. & Hunyadi, B. Modeling Nonlinear Evoked Hemodynamic Responses in Functional Ultrasound. in ICASSPW 2023 - 2023 IEEE International Conference on Acoustics, Speech and Signal Processing Workshops, Proceedings (2023). doi:10.1109/ICASSPW59220.2023.10193541

44. Aguirre, G. K., Zarahn, E. & D’Esposito, M. The variability of human, BOLD hemodynamic responses. Neuroimage 8, (1998).

45. Buxton, R. B., Wong, E. C. & Frank, L. R. Dynamics of blood flow and oxygenation changes during brain activation: The balloon model. Magn. Reson. Med. 39, (1998).

46. Erol, A. et al. Deconvolution of the Functional Ultrasound Response in the Mouse Visual Pathway Using Block-Term Decomposition. Neuroinformatics 21, 247–265 (2022).

47. Erol, A., Van Eyndhoven, S., Koekkoek, S., Kruizinga, P. & Hunyadi, B. Joint Estimation of Hemodynamic Response and Stimulus Function in Functional Ultrasound Using Convolutive Mixtures. in Conference Record - Asilomar Conference on Signals, Systems and Computers 2020-November, (2020).

48. Shelchkova, N. D. et al. Microstimulation of human somatosensory cortex evokes task-dependent, spatially patterned responses in motor cortex. Nat. Commun. 14, (2023).

49. Pons, T. P. & Kaas, J. H. Corticocortical connections of area 2 of somatosensory cortex in macaque monkeys: A correlative anatomical and electrophysiological study. J. Comp. Neurol. 248, (1986).

50. Ghosh, S., Brinkman, C. & Porter, R. A quantitative study of the distribution of neurons projecting to the precentral motor cortex in the monkey (M. fascicularis). J. Comp. Neurol. 259, (1987).

51. Porro, C. A. et al. Primary motor and sensory cortex activation during motor performance and motor imagery: A functional magnetic resonance imaging study. J. Neurosci. 16, (1996).

52. Lugaresi, C., et al. MediaPipe: A Framework for Building Perception Pipelines. ArXiv abs/1906.0, (2019).

53. Hesselmann, V. et al. Discriminating the Cortical Representation Sites of Tongue and Lip Movement by Functional MRI. Brain Topogr. 16, 159–167 (2004).

54. Szameitat, A. J., Shen, S. & Sterr, A. Motor imagery of complex everyday movements. An fMRI study. Neuroimage 34, (2007).

55. Norman, S. L. et al. Single-trial decoding of movement intentions using functional ultrasound neuroimaging. Neuron 109, 554–1566.e4 (2021).

56. Griggs, W. S., et al. Decoding Motor Plans Using a Closed-Loop Ultrasonic Brain-Machine Interface. bioRxiv (2022).

57. Demené, C. et al. Transcranial ultrafast ultrasound localization microscopy of brain vasculature in patients. Nat. Biomed. Eng. 5, 219–228 (2021).

58. Renaudin, N. et al. Functional ultrasound localization microscopy reveals brain-wide neurovascular activity on a microscopic scale. Nat. Methods 19, 1004–1012 (2022).

59. Giacino, J. T., Fins, J. J., Laureys, S. & Schiff, N. D. Disorders of consciousness after acquired brain injury: The state of the science. Nature Reviews Neurology 10, (2014).

60. Edlow, B. L., Claassen, J., Schiff, N. D. & Greer, D. M. Recovery from disorders of consciousness: mechanisms, prognosis and emerging therapies. Nature Reviews Neurology 17, (2021).

61. Schroeder, W., Martin, K. & Lorensen, B. The Visualization Toolkit (VTK). Open Source (2018).

62. Macé, É. et al. Whole-Brain Functional Ultrasound Imaging Reveals Brain Modules for Visuomotor Integration. Neuron 100, 1241–1251.e7 (2018).

63. Guizar-Sicairos, M., Thurman, S. T. & Fienup, J. R. Efficient subpixel image registration algorithms. Opt. Lett. 33, (2008).

