## Supplementary Data for "Mobile Human Brain Imaging using Functional Ultrasound"

**Supplementary Data 1 –** Subject Characteristics

**Table 1 -** Overview of the characteristics of the N=2 subjects included in this study

| Subject | Age Category | Underlying Etiology | Location SBD + PEEK | Handed-ness | Post-operative Neurological Deficits | Baseline Language Assessment | Baseline Cognitive Assessment | Number of measurements |
|---|---|---|---|---|---|---|---|---|
| #1 | 30-35 y | High-Energetic Trauma with multiple cerebral contusions left hemisphere | Hemi-craniectomy Left >4 years in situ | Right | Minimal motor deficits in right arm and leg | Severe Aphasia<br><br>Aphasia Bedside Check: 4/14 Boston Naming Test: 6/60 Token Test: 5.5/36 | Complicated due to aphasia | N = 6 |
| #2 | 35-40 y | Low grade Astrocytoma right insula | Fronto-temporal Right >4 years in situ | Right | None objectified or reported | No deficits | No deficits | N = 2*<br>**Deceased during the course of the study* |

A total of 2 subjects were included in this study, both males in their 30s (**Table 1**). Subject #1 received a left-sided hemicraniectomy and PEEK-implant after high-energetic trauma, causing multiple cerebral contusions and post-operative neurological deficits (**panel A**). Most pronounced was the subject's aphasia, which was considered severe based on the baseline linguistic assessment. Apart from minimal motor deficits in the right arm and leg, which was visible through asymmetries in his gait resulting in a right-sided limp, subject #1 had no other motor-related deficits. Subject #2 received a PEEK-implant in the right-sided frontotemporal region after surgical removal of a low grade astrocytoma in the right insular region. The tumor resection cavity was still clearly visible on MRI-scans (**panel B**). At time of inclusion, the subject was tumor progression-free for multiple years. No cognitive, motor or language deficits were reported or objectified at baseline. Experiments were conducted over a period of 2 years. Subject #1 participated in a total of seven measurements during this period. Subject #2 deceased during the study due to tumor regrowth and participated in two measurements.

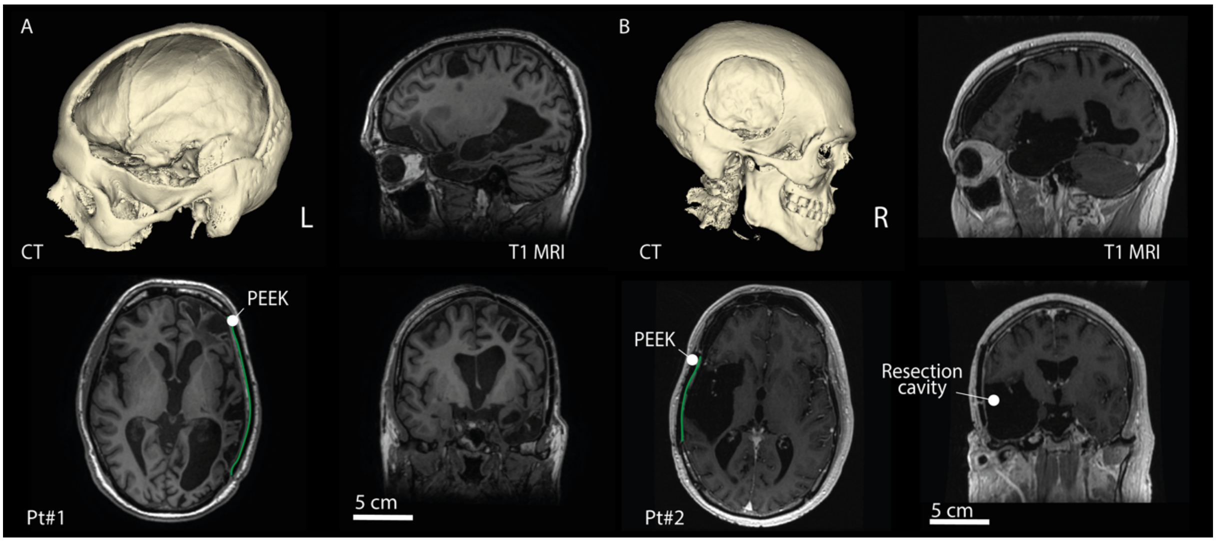

**Supplementary Data 2** – Helmet Fabrication Pipeline

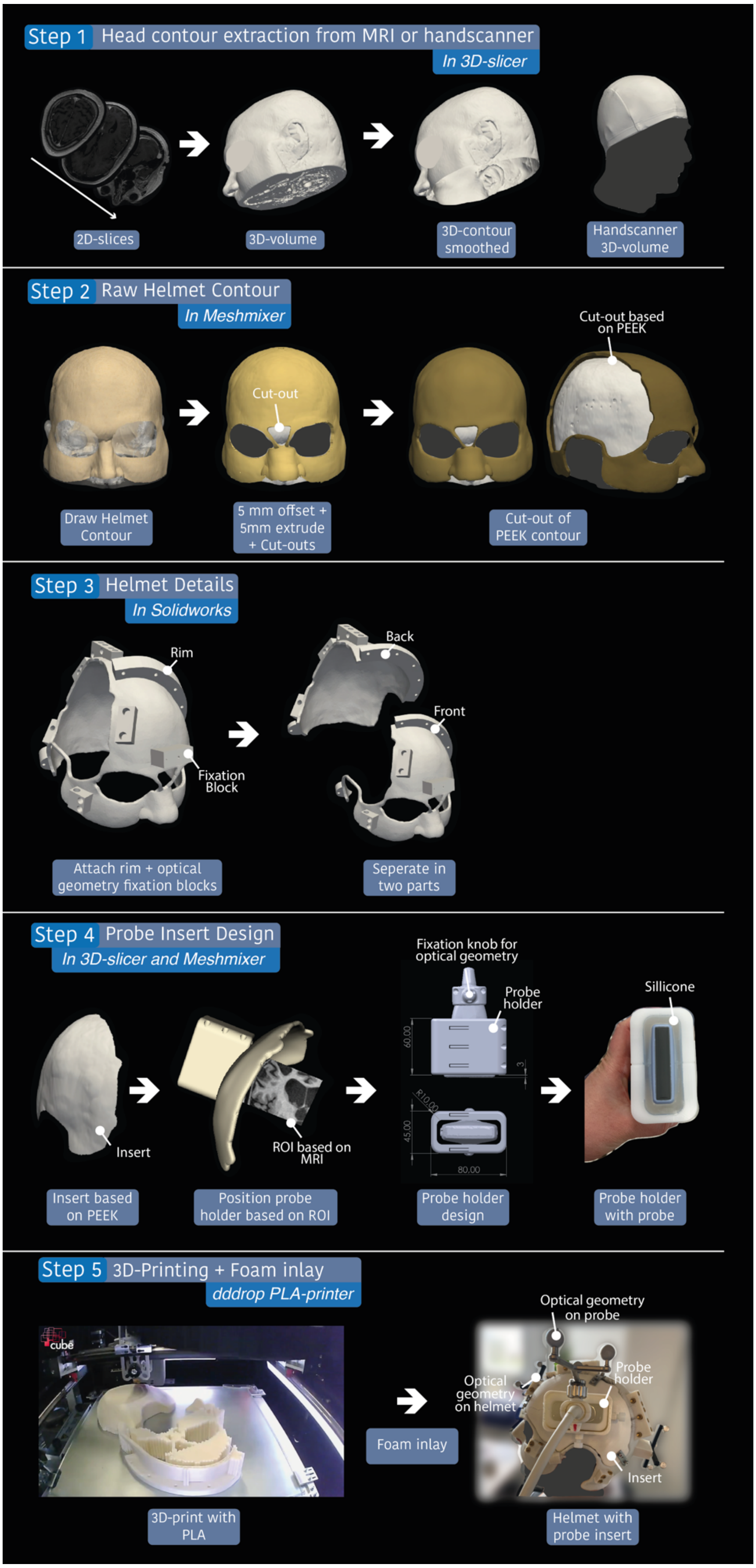

Overview of all the steps involved in the custom helmet design. Step 1 consisted of extracting the contour of the face and head of the subject from clinically available structural MRI datasets of each subject. If no recent MRI-scan was available, an additional 3D surface scan of the face was made using a handheld 3D-scanner and co-registered to the MRI using the landmark registration function in 3D Slicer (this was the case for subject #1). In Step 2 the subject's head contour was smoothed in Meshmixer (Autodesk Inc.), before applying a 5mm offset and 5 mm extrusion to create the base of the fUSi-helmet. Using the same landmark registration function in 3D Slicer, the structural MRI-scan was co-registered to the subject's 3D CT-scan which was made prior to PEEK-production, in order to determine the borders of the defect. A cut-out was made in the helmet with approximately 10 mm margin around the border of the PEEK, which was used as the base for the probe insert, as will be further discussed in Step 4. In Step 3, using SolidWorks (Dassault Systèmes, SolidWorks Corporation) the base of the helmet was expanded with a fixation rim and fixation blocks to mount the necessary geometries for optical tracking (Northern Digital Inc., Canada), see below. The fUSi-helmet was divided in two parts parallel to the fixation rim to allow for printing and fixation on the subject's head. In Step 4 probe inserts were designed based on targeted brain regions of interests (ROIs) for functional tasks using (f)MRI or prior (f)USi-locations. The inserts ensured stability of the probe during functional tasks as well as enable reproducibility of the same 2D-imaging plane across measurements. In order to ensure proper fixation of the ultrasound probe inside this insert, a rectangular probe holder with silicone-lining was molded to the shape of the probe, allowing for fixation of the probe inside the insert. In Step 5, the helmet was 3D-printed in-house using PLA-material (dddrop bv, CAD2M). Finally, the inside of the 3D-printed helmet was covered in foam to ensure subject comfort. Before probe holder containing the probe was positioned in the insert, care was taken to cover the exposed portion of the skull below the insert in a layer of warmed ultrasound-gel, to facilitate optimal acoustic contact.

**Supplementary Data 3** – Optical Tracking Pipeline

Position and orientation of the fUSi-probe and helmet was tracked continuously using an NDI Polaris Vega optical tracking system (SN P9-04539, Northern Digital Inc., Canada), which was configured to track infra-red reflective reference geometries attached to the fUSi-probe and helmet. Custom software was designed to record the tracking information featuring six degrees of freedom (DOF) at an average rate of 20 Hz.

The computation of reference (f)MRI scan slices that corresponded to the ultrasound images required a sequence of transformations on the tracking data. These transformations, provided by the tracking system and through calibration of the tools, facilitated the mapping of a 2D pixel coordinate *(i, j)* in an ultrasound image at a distinct time point *k* to a 3D voxel coordinate *(u, v, w)* in the reference image. The transformation series is described as:

$$\begin{bmatrix} u \\ v \\ w \\ 1 \end{bmatrix} = \mathbf{T}_{\text{img}}^{-1} \cdot \mathbf{T}_{\text{reg}} \cdot \mathbf{T}_{\text{ref},k}^{-1} \cdot \mathbf{T}_{\text{us},k} \cdot \mathbf{T}_{\text{us,local}} \cdot \mathbf{T}_{\text{us,img}} \cdot \begin{bmatrix} i \\ 0 \\ j \\ 1 \end{bmatrix}$$

In this formulation, each **T** embodies a 4x4 affine transformation matrix, encapsulating rotation and translation parameters. Each transformation matrix's role in the sequence is explicated as follows, moving from right to left:

$$\mathbf{T} = \begin{bmatrix} r_{11} & r_{12} & r_{13} & t_x \\ r_{21} & r_{22} & r_{23} & t_y \\ r_{31} & r_{32} & r_{33} & t_z \\ 0 & 0 & 0 & 1 \end{bmatrix}$$

1. $\mathbf{T}_{\text{us,img}}$: Alters pixel coordinates in the ultrasound image into physical positions by scaling them relative to the pixel dimensions and centering the image at the upper middle of the frame.

2. $\mathbf{T}_{\text{us,local}}$: Adjusts the position on the reference geometry attached to the ultrasound probe to align with the center of the ultrasound array.

3. $\mathbf{T}_{\text{us},k}$: Indicates the measured position and orientation of the reference geometry on the fUSi-probe in relation to the tracking camera at time point *k.*

4. $\mathbf{T}_{\text{ref},k}$: Specifies the measured position and orientation of the reference geometry on the fUSi-helmet relative to the tracking camera at time point *k*.

5. $\mathbf{T}_{\text{reg}}$: Denotes the registration transformation mapping the position of the reference geometry on the fUSi-helmet to its position within the anatomical coordinates of the reference (f)MRI image.

6. $\mathbf{T}_{\text{img}}$: Transforms voxel coordinates in the reference image into anatomical coordinates.

To assist in locating specific anatomical regions during hand-held ultrasound scans, we utilized the tracking information in conjunction with software from the Visualization Toolkit (VTK). This combination allowed real-time visualization of the fUSi-probe's position concerning the helmet and several anatomical ROIs, as well as the MRI slice that corresponds to the current ultrasound image (see **Supplementary Data 4**). The tracking data was also saved for later use, facilitating offline 3D reconstruction of the ultrasound path, as demonstrated in **Figure 1B**.

**Supplementary Data 4** – Parallel Datastreams recorded

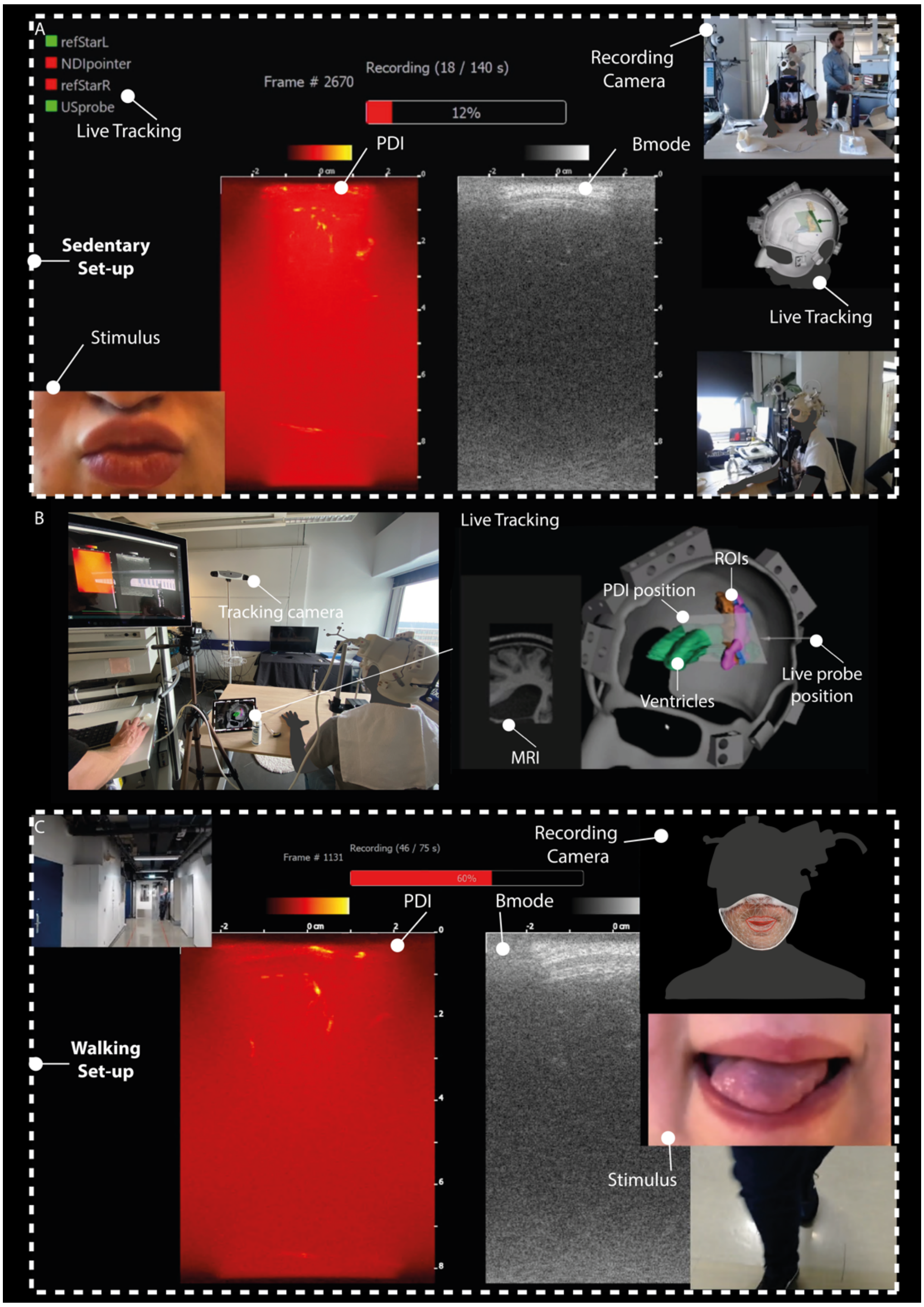


**Photographic images of lips and experimental setting are taken from the authors with explicit permission.*

Screenshot of our GUI demonstrating all the parallel data-streams acquired and stored during our sedentary experiments in the lab (see panel above) and during locomotion (see panel below). The live PDI and Bmode-image as acquired with our custom acquisition unit were recorded in parallel with optical tracking data of the position of the ultrasound probe relative to the patient's helmet and brain anatomy. A live tracking tool was made using VTK[1] to display the probe position relative to the patient anatomy in real-time using previous MRI-scans of the patient (middle panel). Multiple camara-streams were recorded in parallel, as well as the real-time functional stimulus video shown to the patient. This allowed us to study the patient's task performance offline and adjust the stimulus pattern or stimulus delays accordingly.

**Supplementary Data 5** – Ultrasound Distortion Correction

In this study it was possible to visualize functional activity through the PEEK-cranioplasty. However, while the homogeneity of the implant permits imagingcompared to an intact skull it still possesses drawbacks compared to soft-tissue imaging. The interfacial losses caused by transmission through the inset and its higher bulk attenuation both act to decrease the SNR thereby reducing the imaging depth. The impact of these losses on imaging through PMMA has been investigated in a previous study (Rabut et al, 2024)[2]. In addition to these interfacial losses, however, the higher bulk sound speed and complex shape of the implant also induces a lensing effect on the transmitted wave-field. This can be seen in **Figure 1** below which shows a measurement of the acoustic field generated when transmitting a planewave on the clinical transducer both with and without the skull implant. These measurements were taken using a 200 µm needle hydrophone in a custom-built scanning tank. In comparison to free space the shape of the transmitted wavefront is warped, and the time-of-arrival is also shifted.

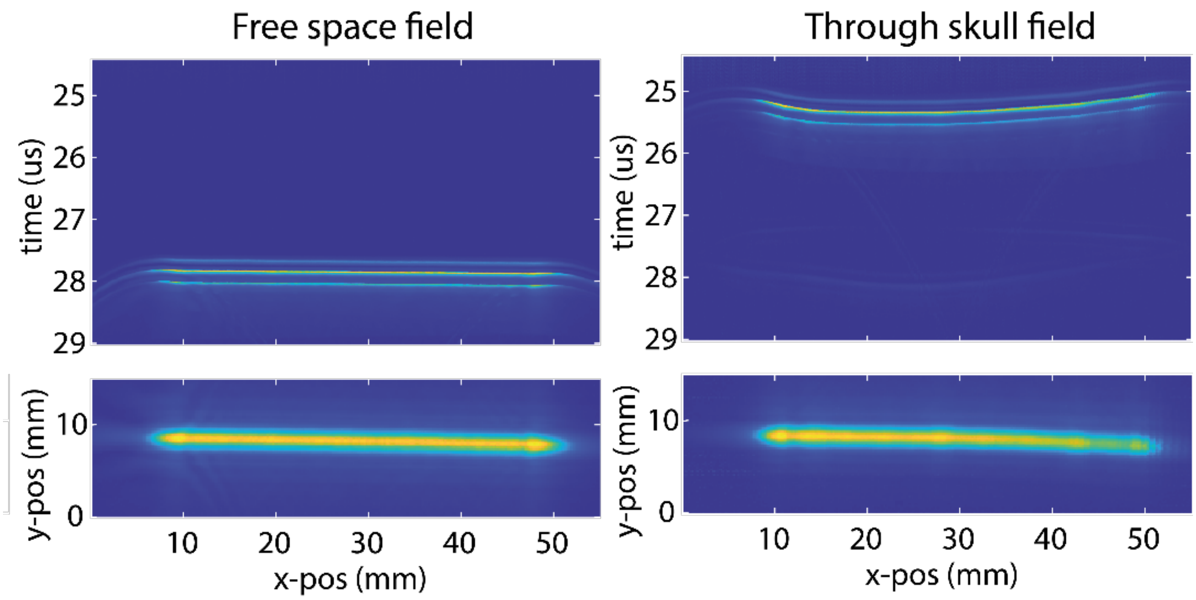


**Figure 1** - (left) Hydrophone scan of acoustic field generated by GE9LD transmitting a normal plane wave in free space. (right) Hydrophone scan of acoustic field generated by GE9LD transmitting a normal plane wave after propagation through skull inset

This distortion of the wavefront and change in time-of-arrival both affect the image reconstruction. First, it makes it necessary to increase the bulk sound speed used in the delay and sum algorithm to compensate for the higher sound speed of the skull implant (2750 ms$^{-1}$ vs 1550 ms$^{-1}$). This increase in the speed of sound can refocus the image, however, only over a limited region as the shift required is proportional to the fraction of the wave path occupied by the skull so changes with depth. Second, by ignoring the lensing of the transmitted wavefront the position and shape of the reconstructed vasculature will be warped compared to its true position. Both effects are illustrated by **Figure 2** which shows reconstructed images of pulse-echo data simulated from a grid of points positioned below a skull implant. Pulse-echo data from the grid of points (**Figure 2a**) was simulated using the k-Wave toolbox, which uses a k-space pseudo-spectral model for time domain simulations of acoustic waves. Images were reconstructed using a delay-and-sum approach using two different values of bulk sound speed - 1700 ms$^{-1}$ (**Figure 2b**) and 1600 ms$^{-1}$ (**Figure 2c**). As the sound speed decreases the depth at which the points are in-focus increases, as the skull occupies a lower-fraction of the wave path. The overall position of the points is also tilted compared to their true position due to the lensing effect of the skull.

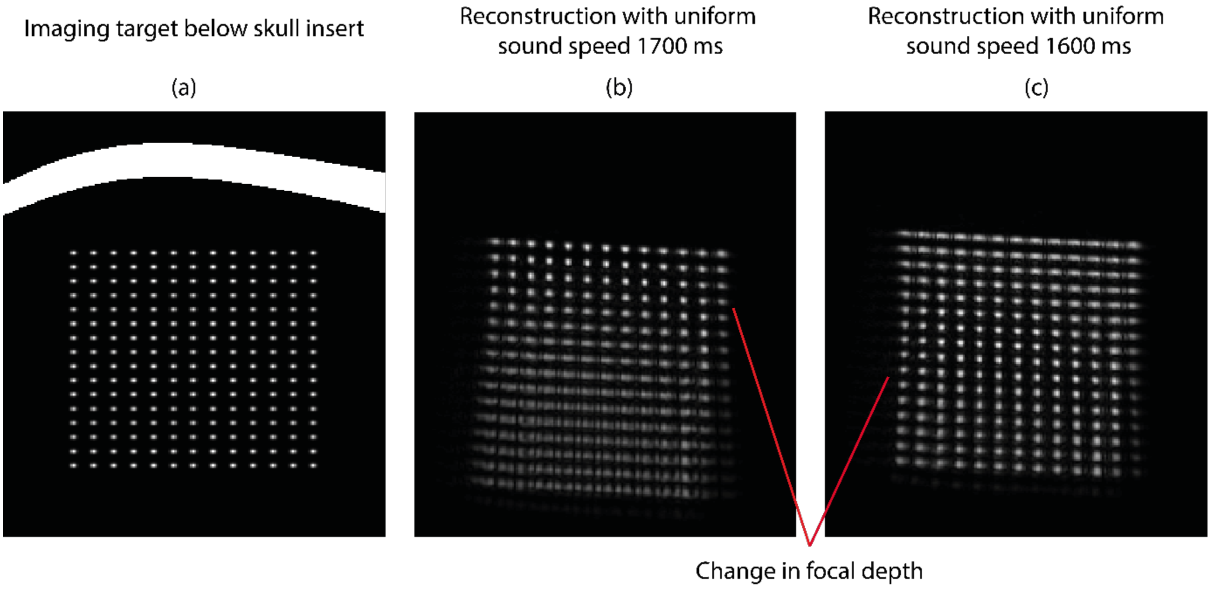


**Figure 2** - Lensing effect of skull insert on reconstructed image. (a) Grid of points behind skull-insert from which pulse-echo data was simulated using the k-Wave toolbox. Increased sound speed of the skull insert can be compensated by increasing the sound speed, however, this results in an image that is only partially in focus. The underlying image structure is also warped.

We investigated whether it was possible to compensate for this by using a model-based approach for image reconstruction. For this model-based reconstruction approach we assume our data $y$ can be linearly related to our image $x$ via a matrix vector multiplication $y = Ax$, where the matrix $A$ contains the pulse-echo impulse response for each pixel in the imaging medium. This framing allows for a variety of different solvers to be used for the image reconstruction.

We then tried to approximate a matrix $A$ that incorporated the effect of the implant using an image-guided approach. First, we reconstructed an image using a model $A$ calculated assuming a homogeneous sound speed of 1550 ms$^{-1}$ matching soft tissue. (**Figure 3a**). To calculate $A$ we simulated the forward field for each element of the linear array using k-Wave then constructed the pulse-echo impulse response assuming linearity and reciprocity using an approach reported previously (Brown et al, 2024)[3]. The image was reconstructed using matched filter (e.g., $x = A^H y$). This generated an image that was largely out of focus (**Figure 3b**), however, could be used to manually segment the top-surface of the skull implant. Using this segmented surface we generated a new sound speed map for our simulation domain comprising a half-space of soft-tissue and the skull insert (**Figure 3c**). The model was then recomputed in this half-space using k-Wave to model the wave propagation and a second B-mode image was reconstructed from which the bottom surface of the skull implant could be manually segmented (**Figure 3d**). With the segmented bottom surface, we were able to generate a final map of acoustic properties containing the full skull-bone implant shape. The model was again updated using k-Wave. We found that when using this final model the resulting B-mode image was visually sharper throughout the field-of-view having compensated for the lensing of the skull insert (**Figure 3f**).

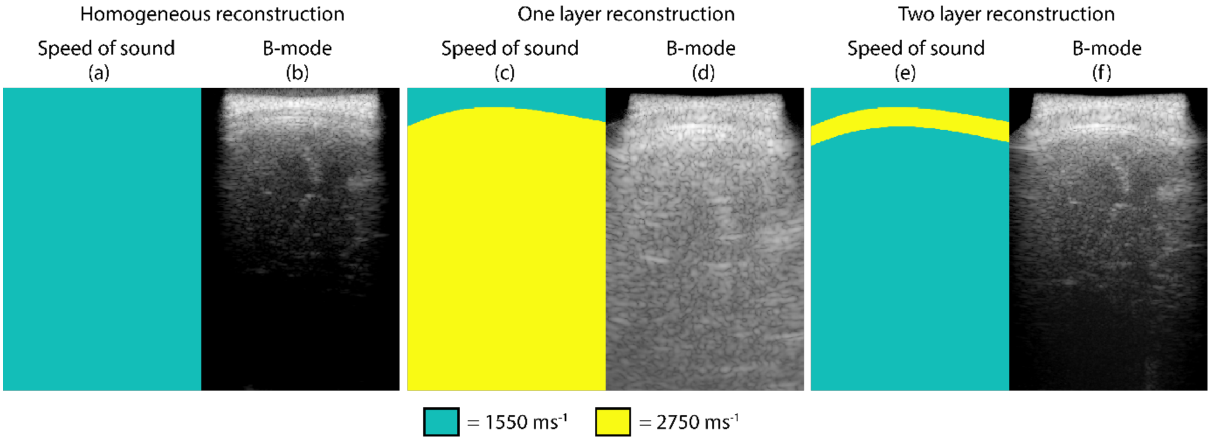


**Figure 3** - Image-guided approach used to correct for the lensing effect of the skull insert. (a-b) sound speed map and resulting B-mode image used for segmenting the first skull layer. (c-d) sound speed map and resulting B-mode image used for segmenting the second skull layer. (e-f) final sound speed map and B-mode image that corrects for the lensing effect of the skull insert.

We recomputed one set of PDIs using the updated model using 200 frames of raw RF data. For comparison we also reconstructed a PDI from the same data using a bulk sound speed of 1700 ms$^{-1}$. It can be seen in **Figure 4** that the resulting visual correspondence is significantly improved.

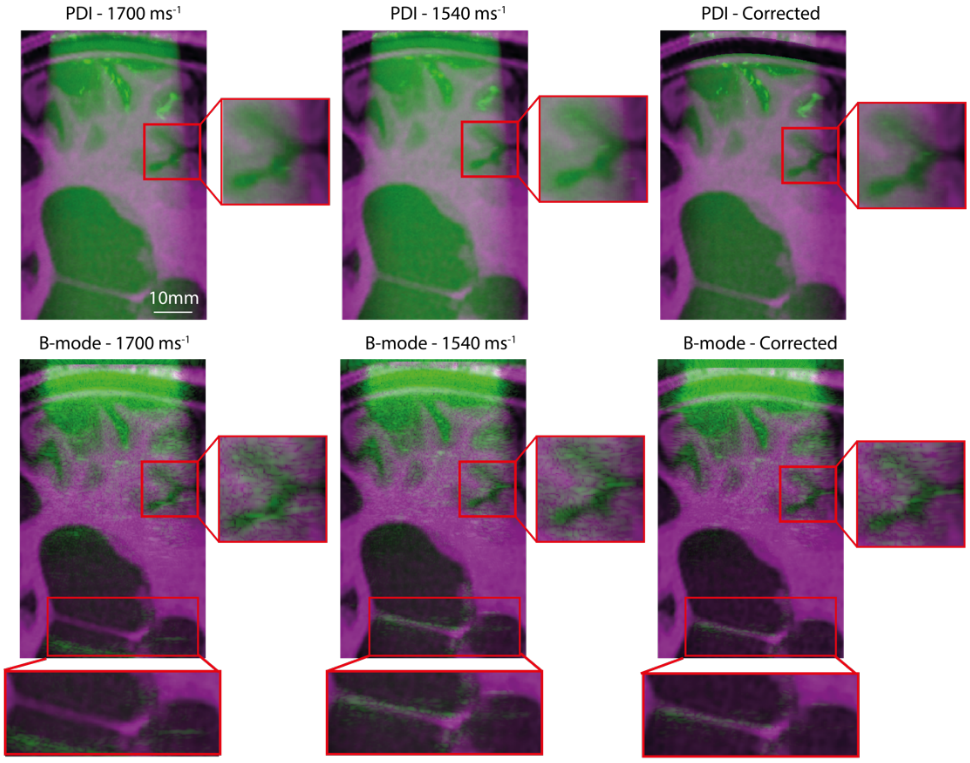


**Figure 4** – Improved visual correspondence between the PDI corrected using our proposed methods (most right panel), versus the PDI reconstructed using a bulk sound speed of 1700 ms$^{-1}$ or 1540 ms$^{-1}$ (left and middle panel, respectively).

**Supplementary Data 6** – Hemodynamic response function estimation

The temporal fluctuations of the functional signal *y(t)* recorded during task is commonly modelled as the response of linear time-invariant (system) to the task time course *u(t).* The impulse response of the LTI system is known as the hemodynamic response function (HRF) $h^{(1)}(t)$. Considering a certain baseline fUSi signal $h^{(0)}$ , the functional signal time series can be written as:

$$y(t) \approx h^{(0)} + \sum_{t_1=0}^{T-1} h^{(1)}(t_1)u(t-t_1) \quad (1)$$

Based on the above mathematical model, and using the fUSi measurements as well as the known stimulus time course, the parameters h(0) and h(1)(t) can be estimated. To ensure that the estimated parameters produce a plausible HRF shape, the kernel h(1)(t) can be expanded as the linear combination of L temporal basis functions b_i(k), k=1,..,L. The basis functions were chosen in the form of gamma functions:

$$b(k;\boldsymbol{\theta}) = \theta_1\left(\Gamma(\theta_2)\right)^{-1}{\theta_3}^{\theta_2}k^{\theta_2-1}e^{-\theta_3 k} \quad (2)$$

The parameters were varied to account for different possible peak delays, equidistant between from the stimulus onset. The basis functions for L=10 are depicted in **Figure 1**. The detailed estimation methodology is described in Kotti et al. (2023)[4], with the difference that in this work the kernel was not restricted to be positive only.

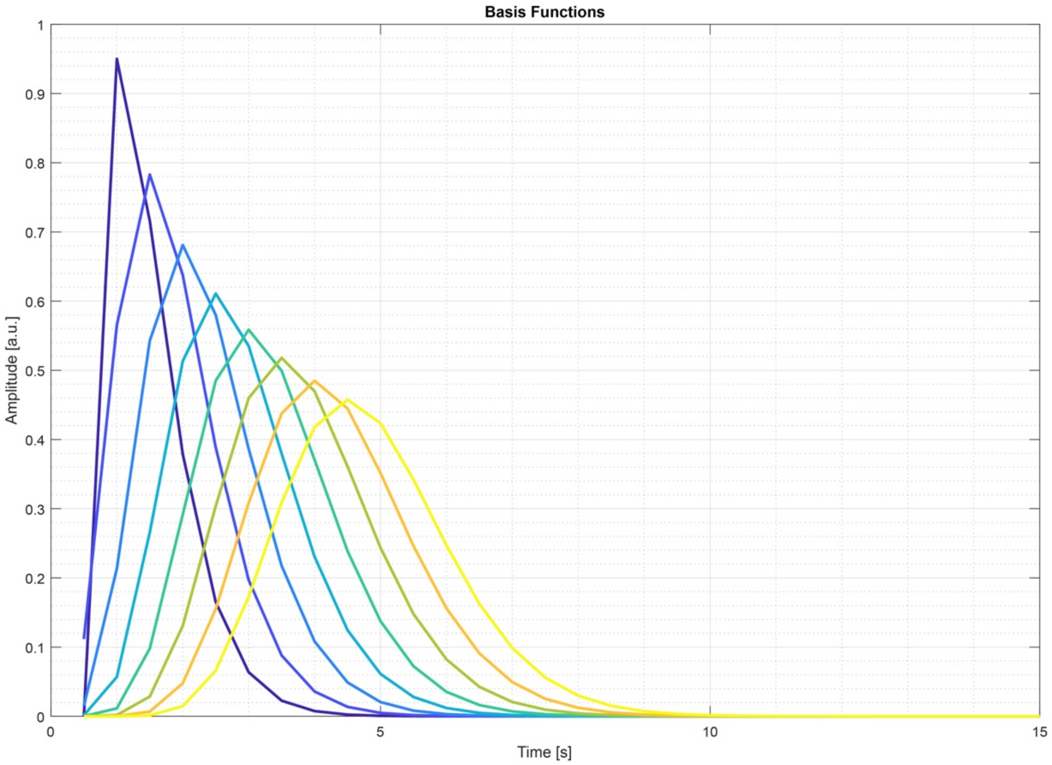


**Figure 3** - Basis functions with L=8

We used a total of 4, 55s long fUSi recordings for HRF estimation. The optimal number of basis functions L as well as the l1-regularization coefficient λ controlling the number of basis functions with non-zero weight were determined in a 4-fold cross-validation setting. Namely, we concatenated all but one measurements for estimating the HRF, then convolved the tracking time course with the estimated HRF to predict the left-out fUSi time series using equation (1). To evaluate the quality of the estimation, the PCC and mean squared error between the predicted and measured fUSi were calculated. Based on this cross-validation the values L=10 and λ=0.1 were chosen. After establishing these optimal values, all 4 measuremetns were concatenated to estimate the final HRF to be used in our further analysis. The resulting HRF is shown in **Figure 2** (dotted green line) overlayed on estimated HRFs from each fold in the cross-validation (thin lines).

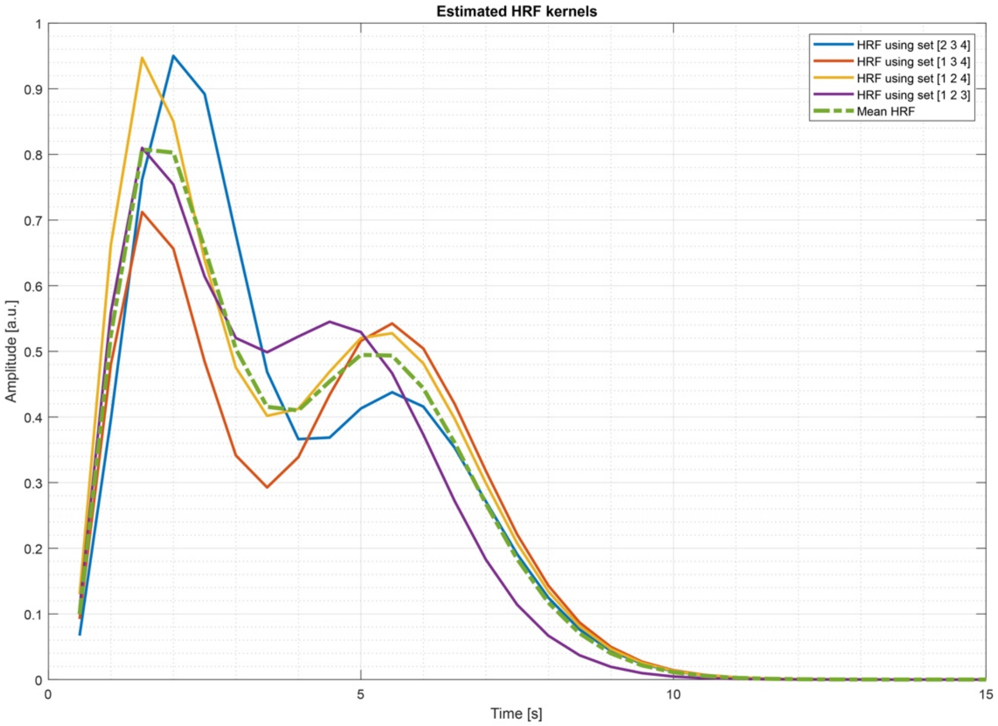


**Figure 4** - Estimated HRF based on all measurements (dotted green) overlayed on estimated HRF from each fold in the cross-validation (thin lines).

To illustrate the reliability of the estimated fUSi time courses using the above HRF, we applied this HRF in an independent set of measurements. In **Figure 3** below, the raw fUSi time courses are shown in blue, the tracking time course (shifted with 3.5s) in black and the estimated fUSi time course in orange. The estimated fUSi time series achieves higher correlations with the measured time course compared to the optimally delayed tracking signal (r_hrf and r_raw, respectively, shown above the time traces).

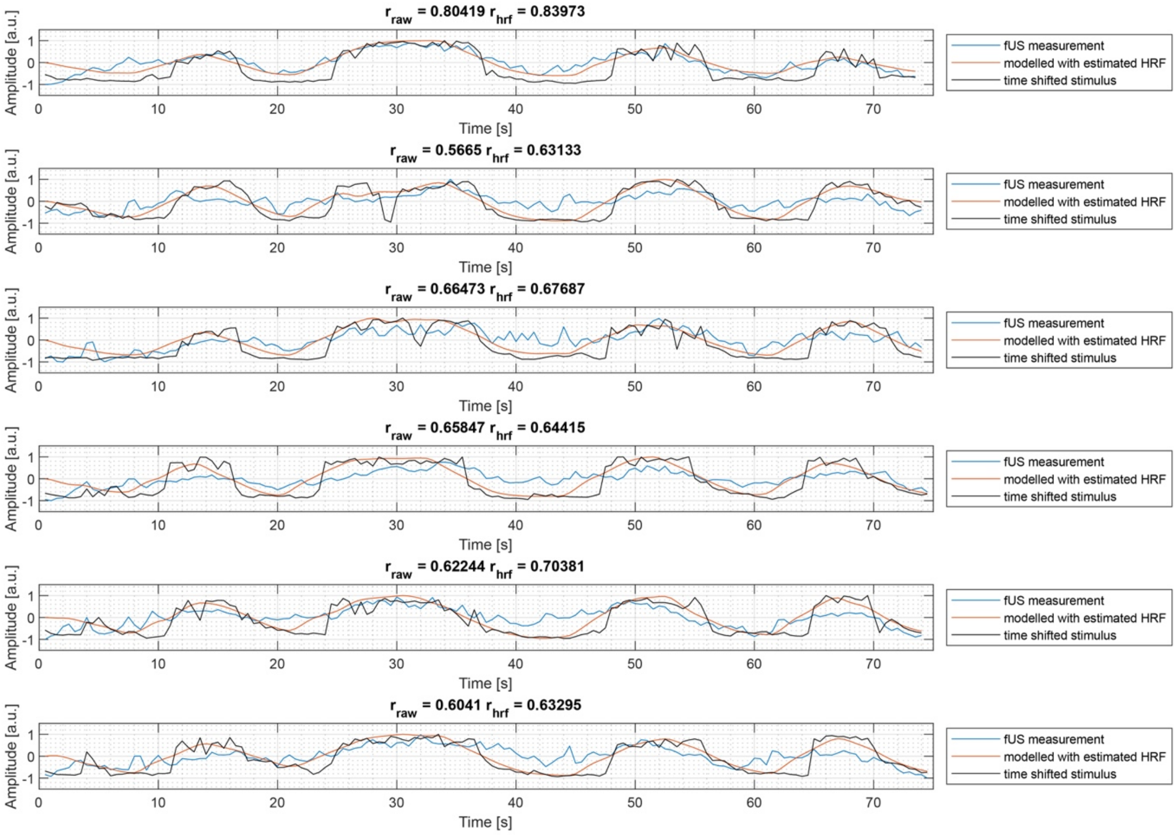


**Figure 5** - Estimated fUSi time courses (orange) using the estimated HRF shown on Figure 2. For comparison, measured fUSi time courses are shown in blue and the tracking time course (delayed with an optimal delay of 3.5s) is shown in black. Correlations between the fUSi measurements and the estimated fUSi time courses (r_hrf) are higher than correlations between the measurements and the optimally delayed tracking time courses (r_raw).

### **Supplementary Data 7 –** Overview of functional tasks used during fUSi and fMRI

| Figure Panel | Modality | Task Content | Task Pattern | ON-task | OFF-task |
|---|---|---|---|---|---|
| 2E | fMRI | Lip Pouting (motor, video-guided) | 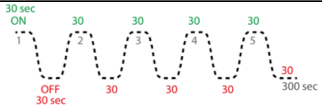 | 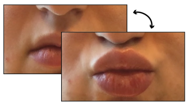 | Rust. U hoeft niks te doen. |
| 2D | fUSi | Lip Pouting (motor, video-guided) | 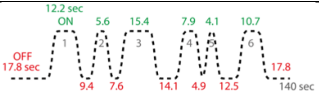 | 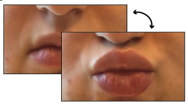 | Rust. U hoeft niks te doen. |
| 2H | fUSi | Lip Sensory (brushing by experimenter) | 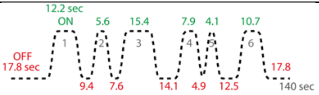 | Lip brushing | |
| 3C | fUSi | Lip Pouting (motor, video-guided) | 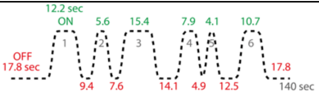 | 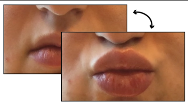 | Rust. U hoeft niks te doen. |
| 4A | fUSi | Lip Sensory (brushing by experimenter) | 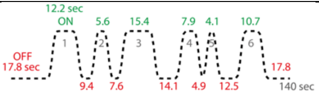 | Lip brushing | |
| 4B | fUSi | Lip Sensory Forehead in off (brushing by experimenter) | 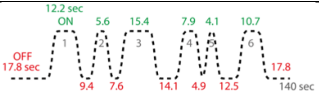 | Lip brushing | Forehead brushing |
| 4C | fUSi | Lip Sensory Ear in off (brushing by experimenter) | 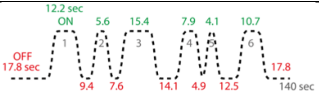 | Lip brushing | Ear brushing |
| 4D | fUSi | Lip Sensory Hand in off (brushing by experimenter) | 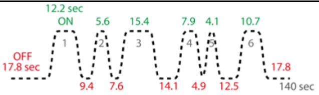 | Lip brushing | Hand brushing |
| 4E | fUSi | Forehead Sensory (brushing by experimenter) | 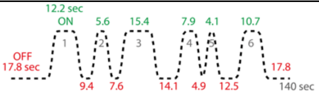 | Forehead brushing | |
| 4F | fUSi | Lip Pouting (motor, video-guided) | 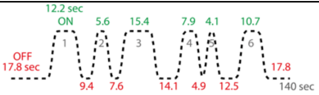 | 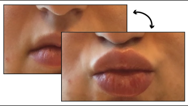 | Rust. U hoeft niks te doen. |
| 4G | fUSi | Lip Sensory Imagined (Video-guided) | 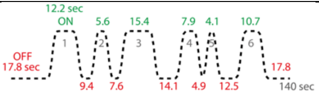 | 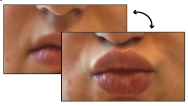 | Rust. U hoeft niks te doen. |
| 4J | fUSi | Lip Sensory (continuous brushing by experimenter) | 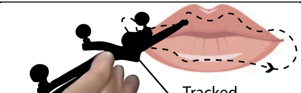 Tracked Lip brush | *Continuous* | *Continuous* |
| 5D | fUSi | Lip Licking (video-guided) | 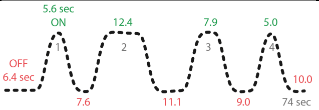 | 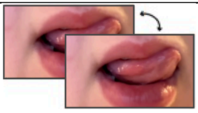 | Rust. U hoeft niks te doen. |
| 5F | fUSi | Lip Licking (video-guided) | 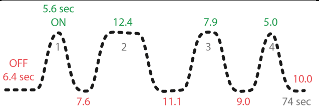 | 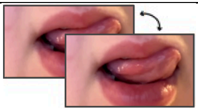 | Rust. U hoeft niks te doen. |
| 5G | fUSi | Lip Licking (audio-guided) | 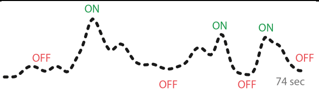 *Depending on the tracked lip trace* | 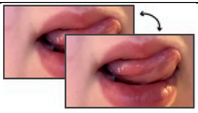 | Rust. U hoeft niks te doen. |

| | | | | | |
|---|---|---|---|---|---|
| 5H | fUSi | Lip Licking (imagined) | 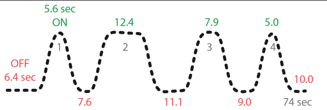 | 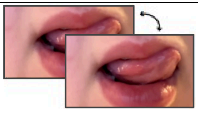 | 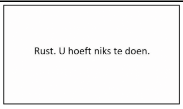 |
| 5I | fUSi | Finger Tapping (imagined) | 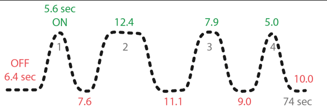 | 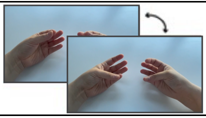 | 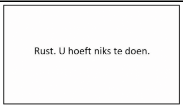 |
| 5J | fUSi | Data from 5G, 5H and 5I combined in one scatterplot | *See above* | *See above* | *See above* |

**Photographic images of lips and hands are taken from one of the authors with explicit permission.*

**Supplementary Data 8** – Average motion during walking tasks

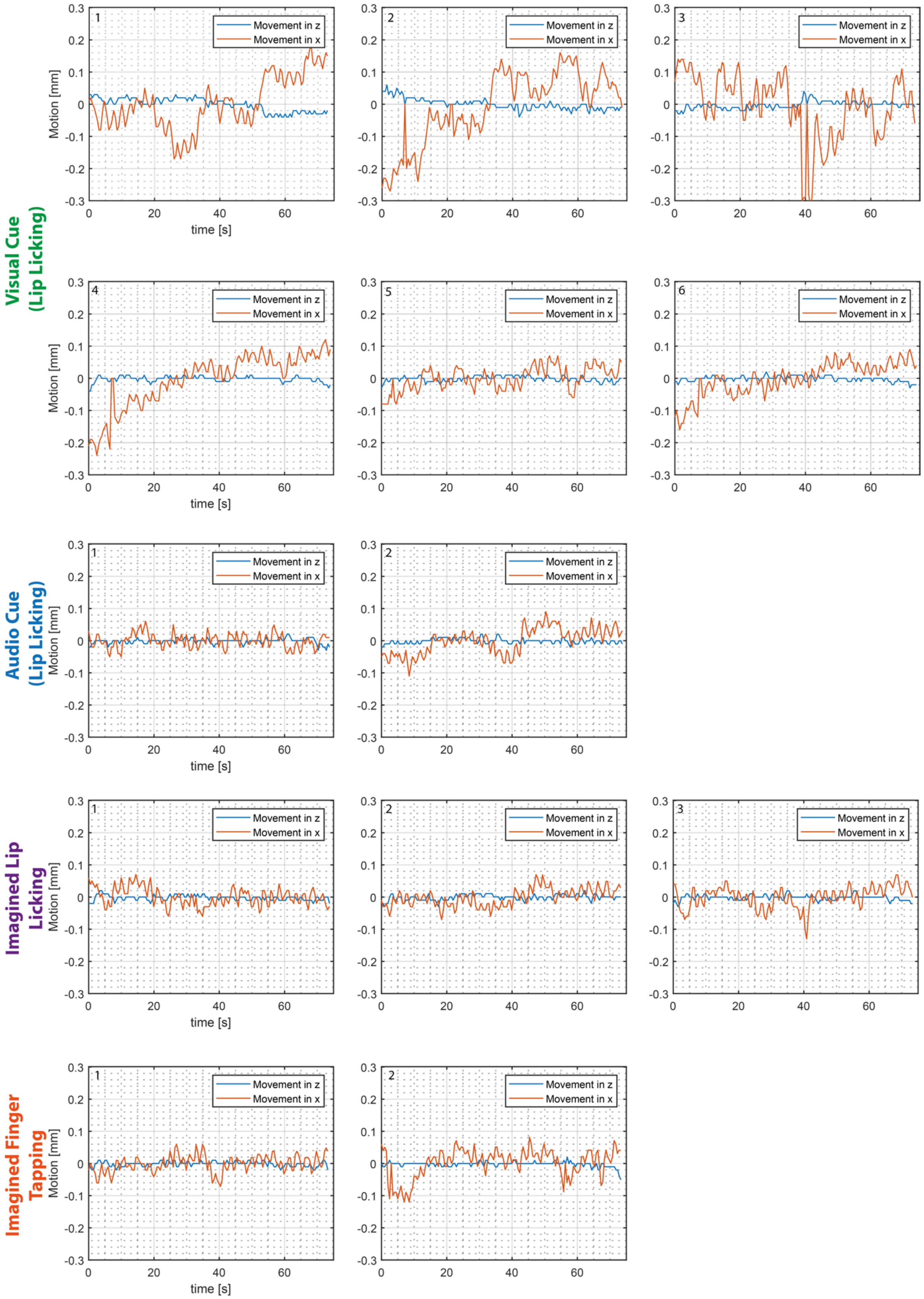

**Supplementary Data 9** – Conventional ultrasound images through PEEK

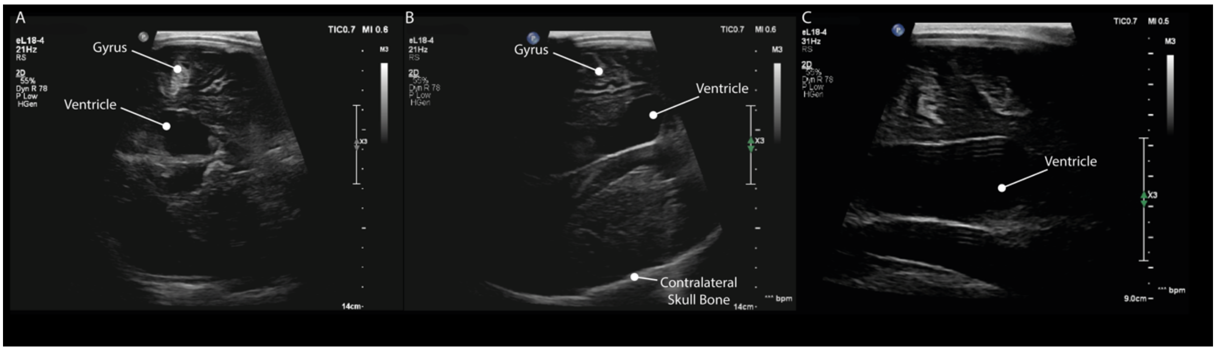


In order to put our fUS-images in perspective, we made images through PEEK in pt.#1 using a commercial-grade ultrasound machine (Philips EPIQ Elite), interfaced with an eL18-4 linear array (Philips, 1920 elements, 21 MHz center frequency). The sonotransparancy of the PEEK is confirmed again with the Bmode images we were able to make, showing even the contralateral skull bone in view (see panel B).

**Supplementary Data 10** – Datasheet

| Figure Panel | Subject | Content | Recording ID (internal code) | Date of data collection | Recording Duration |
|---|---|---|---|---|---|
| 1C | #1 | Free-hand sweep over cranioplasty (vascular data only) | 522 | 10-05-2023 | 49 sec |
| 2B | #1 | Motor lip pouting task (ON (lip pouting) – OFF (nothing)) | 546<br>**Ne = 600* | 10-05-2023 | 140 sec |
| 2C | #1 | Motor lip pouting task (ON (lip pouting) – OFF (nothing)) | 444 \| 546\|<br>20240902T151431<br>Ne = 800 | 07-04-2022\|<br>10-05-2023\|<br>02-09-2024 | 140 sec |
| 2D | #1 | Motor lip pouting task (ON (lip pouting) – OFF (nothing)) | 444 \| 546\|<br>20240902T151431 | 07-04-2022\|<br>10-05-2023\|<br>02-09-2024 | 140 sec |
| 2G | #1 | Sensory lip brushing task (ON (lip brushing) – OFF (nothing)) | 531\|551 | 10-05-2023\|<br>10-05-2023 | 140 sec |
| 2H | #1 | Sensory lip brushing task (ON (lip brushing) – OFF (nothing)) | 531\|551 | 10-05-2023\|<br>10-05-2023 | 140 sec |
| 3B | #2 | Motor lip pouting task (ON (lip pouting) – OFF (nothing)) | 467 | 13-04-2022 | 140 sec |
| 3C | #2 | Motor lip pouting task (ON (lip pouting) – OFF (nothing)) | 467\|507 | 13-04-2022 \|<br>06-07-2022 | 140 sec |
| 4A | #1 | Sensory lip brushing task (ON (lip brushing) – OFF (nothing)) | 531 | 10-05-2023 | 140 sec |
| 4B | #1 | Sensory lip brushing task (ON (lip brushing) – OFF (forehead)) | 537 | 10-05-2023 | 140 sec |
| 4C | #1 | Sensory lip brushing task (ON (lip brushing) – OFF (ear)) | 544 | 10-05-2023 | 140 sec |
| 4D | #1 | Sensory lip brushing task (ON (lip brushing) – OFF (hand)) | 533 | 10-05-2023 | 140 sec |
| 4E | #1 | Sensory lip brushing task (ON (forehead) – OFF (nothing)) | 540 | 10-05-2023 | 140 sec |
| 4F | #1 | Motor lip pouting task (ON (lip pouting) – OFF (nothing)) | 546 | 10-05-2023 | 140 sec |
| 4G | #1 | Imagined Lip Licking (look at video cue with ON-OFF lip licking without motor movement) | 529 | 10-05-2023 | 140 sec |
| 4J | #1 | Lip Sensory<br>(continuous brushing by experimenter) | 621 | 13-12-2023 | 178 sec |
| 4K | #1 | Lip Sensory<br>(continuous brushing by experimenter) | 621 | 13-12-2023 | 178 sec |
| 5C | #1 | Lip Licking while walking (video-guided, ON (lip licking), OFF (nothing)) | 640 | 20-12-2023 | 75 sec |
| 5D | #1 | Lip Licking while walking (video-guided, ON (lip licking), OFF (nothing)) | 640 | 20-12-2023 | 75 sec |
| 5E | #1 | Lip Licking while walking (video-guided, ON (lip licking), OFF (nothing)) | 639 \| 640 \| 641\| 642 \| 643 \| 644 | 20-12-2023 | 75 sec |
| 5F | #1 | Lip Licking while walking (video-guided, ON (lip licking), OFF (nothing)) | 639 \| 640 \| 641\| 642 \| 643 \| 644 | 20-12-2023 | 75 sec |
| 5G | #1 | Lip Licking while walking (audio-guided, ON (lip licking), OFF (nothing)) | 645 \| 646 | 20-12-2023 | 75 sec |
| 5H | #1 | Imagined Lip Licking while walking (look at video cue with | 647 \| 648 \| 649 | 20-12-2023 | 75 sec |

| | | | | | |
|---|---|---|---|---|---|
| | | ON-OFF lip licking without motor movement) | | | |
| 5I | #1 | Imagined finger tapping while walking (look at video cue with ON-OFF finger tapping without motor movement) | 650 \| 651 | 20-12-2023 | 75 sec |
| 5J | #1 | *See above* | All the datasets of panels 5G-I combined | *See above* | *See above* |
| Suppl. Data 5 | #1 | Motor lip pouting task (ON (lip pouting) – OFF (nothing)) | 546 | 10-05-2023 | 140 sec |
| Suppl. Data 6 | #1 | Lip Licking while walking (video-guided, ON (lip licking), OFF (nothing)) (prior dataset recorded for training purposes) | 607 \|608 \| 609 \| 610<br><br>639 \| 640 \| 641\| 642 \| 643 \| 644 | 13-12-2023<br><br>20-12-2023 | 56 sec<br><br>75 sec |
| Suppl. Data 8 | #1 | *See above* | All the datasets of panels 5G-I combined | *See above* | *See above* |

**Ne = Ensemble Size. If no Ne is defined, the standard Ne = 800 was applied.*

**References in Supplementary Data**


1. Schroeder, W., Martin, K. & Lorensen, B. The Visualization Toolkit (VTK). *Open Source* (2018).
2. Rabut, C. *et al.* Functional ultrasound imaging of human brain activity through an acoustically transparent cranial window. *Sci. Transl. Med.* **16**, eadj3143 (2024).
3. Brown, M. D. *et al.* Four-dimensional computational ultrasound imaging of brain hemodynamics. *Sci. Adv.* **10**, (2024).
4. Kotti, S. E., Erol, A. & Hunyadi, B. Modeling Nonlinear Evoked Hemodynamic Responses in Functional Ultrasound. in *ICASSPW 2023 - 2023 IEEE International Conference on Acoustics, Speech and Signal Processing Workshops, Proceedings* (2023).
